# Iron acquisition system of *Sphingobium* sp. strain SYK-6, a degrader of lignin-derived aromatic compounds

**DOI:** 10.1101/2020.03.26.005173

**Authors:** Masaya Fujita, Taichi Sakumoto, Kenta Tanatani, Hong Yang Yu, Kosuke Mori, Naofumi Kamimura, Eiji Masai

## Abstract

Iron, an essential element for all organisms, acts as a cofactor of enzymes in bacterial degradation of recalcitrant aromatic compounds. The bacterial family, Sphingomonadaceae comprises various degraders of recalcitrant aromatic compounds; however, little is known about their iron acquisition system. Here, we investigated the iron acquisition system in a model bacterium capable of degrading lignin-derived aromatics, *Sphingobium* sp. strain SYK-6. Analyses of SYK-6 mutants revealed that FiuA (SLG_34550), a TonB-dependent receptor (TBDR), was the major outer membrane iron transporter. Three other TBDRs encoded by SLG_04340, SLG_04380, and SLG_10860 also participated in iron uptake, and *tonB2* (SLG_34550), one of the six *tonB* comprising the Ton complex which enables TBDR-mediated transport was critical for iron uptake. The ferrous iron transporter FeoB (SLG_36840) played an important role in iron uptake across the inner membrane. The promoter activities of most of the iron uptake genes were induced under iron-limited conditions, and their regulation is controlled by SLG_29410 encoding the ferric uptake regulator, Fur. Although *feoB*, among all the iron uptake genes identified is highly conserved in Sphingomonad strains, the outer membrane transporters seem to be diversified. Elucidation of the iron acquisition system promises better understanding of the bacterial degradation mechanisms of aromatic compounds.

## Introduction

Iron is an essential nutrient utilised as a cofactor for enzymes that control various life phenomena such as respiration, dissimilation, and stress response^1^. Iron exists mainly in ferrous and ferric forms. Ferrous iron is soluble and highly bioavailable; however, the predominant form of iron in the environment is an insoluble ferric form^2^. Ferric iron in soil and water is solubilised by forming complexes with organic ligands such as siderophores, humic acid, lipids, proteins, and polysaccharides^3,4^. Many bacteria secrete specific high-affinity siderophores which form complexes with ferric iron and then uptake these ferric-siderophore complexes to enhance iron acquisition over competitors^5–7^. Besides, pathogenic bacteria can acquire haem and transferrin specifically from their host cells^8,9^.

Gram-negative bacteria need to transport iron through both the outer and inner membranes. TonB-dependent receptors (TBDRs) mediate transport of ferric complexes (e.g. siderophore, haem, and transferrin) across the outer membrane^1,10,11^. TBDRs utilize energy derived from the proton motive force transmitted by TonB-ExbB-ExbD complex (Ton complex) localised in the inner membrane (Ton system)^12^. Beside siderophores, the Ton system is involved in the uptake of vitamin B12, saccharide, aromatic compounds, and metals such as nickel, copper, and lanthanoid^13–18^. The uptake of the ferric complex across the inner membrane is mainly achieved by ATP-binding cassette (ABC) transporters^11^. A major facilitator superfamily (MFS) transporter, FptX, is also known to mediate the ferric-siderophore (pyochelin) uptake in *Pseudomonas aeruginosa*^19^. The ferric iron transported into the cytoplasm is eventually reduced to ferrous iron.

The outer membrane transport of free ferrous and ferric irons is thought to be mediated by porins^11,20^. A recent study has suggested that TBDRs of *Synechocystis* sp. PCC 6803 mediate the uptake of free iron, thereby not restricting iron uptake to only ferric-siderophore complexes^21^. The transport of ferrous iron across the inner membrane is mainly facilitated by Feo and divalent metal ion transporters such as MntH and ZupT^20^. The Feo system encoded by *feoABC* is considered the primary uptake system for ferrous iron in both pathogenic and non-pathogenic bacteria^20^. FeoB is a permease that utilizes energy acquired by its N-terminus GTPase domain and transports ferrous iron. FeoA and FeoC are accessory proteins important for FeoB multimer formation, and essential for growth utilizing ferrous iron in *Vibrio cholerae*^22,23^. However, *feoC* is conserved in only gammaproteobacteria, and the organisation of the *feo* operon varies among bacteria^20^.

The transcription of most of the genes involved in iron uptake and metabolism is regulated by the ferric uptake regulator (Fur)^24,25^. Excess ferrous iron results in the binding of the ferrous iron-Fur complex to the Fur box in the promoter regions to repress their transcription.

Until now, iron acquisition by proteobacteria has been mainly investigated in pathogenic bacteria^11^. To the best of our knowledge, reports regarding the transporters involved in iron acquisition by alphaproteobacteria are limited to Rhizobiales and Caulobacterales^19,26,27^. The family Sphingomonadaceae in alphaproteobacteria comprises of many unique strains such as *Sphingobium* sp. SYK-6, *Sphingomonas wittichii* RW1, *Novosphingobium pentaromativorans* US6-1, and *Sphingobium chlorophenolicum* L-1, capable of degrading certain recalcitrant aromatic compounds such as lignin-derived aromatic compounds, dibenzo-*p*-dioxin, polycyclic aromatic hydrocarbons, and pentachlorophenol, respectively^28–31^. These strains are valuable for bioremediation and the production of industrially useful chemicals from biomass^28,29,32^. Iron, an essential factor in the bacterial degradation of aromatic compounds is located in the active centre of *O*-demethylases, aromatic-ring-hydroxylating oxygenases, and ring-cleavage enzymes^33,34^. *Sphingobium* sp. SYK-6 produces 2-pyrone-4,6-dicarboxylate (PDC), a promising platform chemical that enables the synthesis of functional polymers, during the degradation of lignin-derived aromatics, thereby indicating that the SYK-6 catabolic system is useful to lignin valorisation^35–37^. Iron also has essential roles in the catabolism of lignin-derived aromatic compounds as exemplified by the presence of ferrous iron in the active centres of protocatechuate 4,5-dioxygenase (LigAB), gallate dioxygenase (DesB), 3-*O*-methylgallate dioxygenase (DesZ), and a ring cleavage dioxygenase of an intermediate metabolite of 5,5′-dehydrodivanillate (DDVA) (LigZ)^38–41^. Ferrous iron is also contained in the active centre and an iron-sulphur cluster of the oxygenase (LigXa) and the ferredoxin (LigXc) components of the multicomponent DDVA *O*-demethylase, respectively^42^. Analysis of the outer membrane transporters of lignin-derived aromatic compounds in SYK-6 has indicated that *ddvT*, one of the 74 TBDR genes, encodes the DDVA outer membrane transporter, and *tonB1* is involved in this transport among the six *tonB*^15^. On the other hand, disruption of *tonB2* was seen to affect the growth of SYK-6 and decrease the activity of ferrous iron-requiring DDVA *O*-demethylase, thereby suggesting that *tonB2* plays a role in the iron acquisition process.

In this study, we identified the SYK-6 transporters mainly involved in the uptake of iron across the outer and inner membranes through the analyses of mutants of the candidate iron uptake genes, their promoter activities in response to iron, and the binding of Fur to their promoter regions to gain insight into the iron acquisition system of Sphingomonadaceae. Such detailed insight into the iron acquisition system of these recalcitrant compound degrading bacteria could have immense implications in enhancement of biodegradability of certain compounds which could extrapolate to improvement of environmental conditions.

## Results and Discussion

### Identification of *tonB* involved in iron uptake

We examined the capacity of the wild type and *tonB2-6* mutants (a *tonB1* mutant was unable to be obtained) to grow on vanillate (VA), protocatechuate (PCA), and SEMP (10 mM sucrose, 10 mM glutamate, 20 mg l^−1^ methionine, and 10 mM proline)^43^ in the presence (the iron-limited condition) and absence (the iron-replete condition) of 100 μM 2,2′-dipyridyl (DIP) to identify the particular *tonB* involved in the iron uptake among the six *tonB*. While Δ*tonB2* cells only showed growth retardation when grown on VA and SEMP under iron-replete conditions (Fig. 1), under iron-limited conditions, Δ*tonB2* cells showed further growth retardation and almost lost the capacity to grow on VA. However, the growth characteristic of Δ*tonB3-6* was mostly the same as that of the wild type. Although under iron-replete conditions, the growth of Δ*tonB2-6* cells on PCA matched that of the wild type, iron limiting conditions showed slight growth retardation of Δ*tonB2* cells on PCA. The growth of Δ*tonB2* cells on VA, PCA, and SEMP under iron-limited conditions was recovered by the introduction of a *tonB2*-carrying plasmid, indicating that the growth retardation described above was caused by the disruption of *tonB2* (Fig. S1).

**Figure 1.**
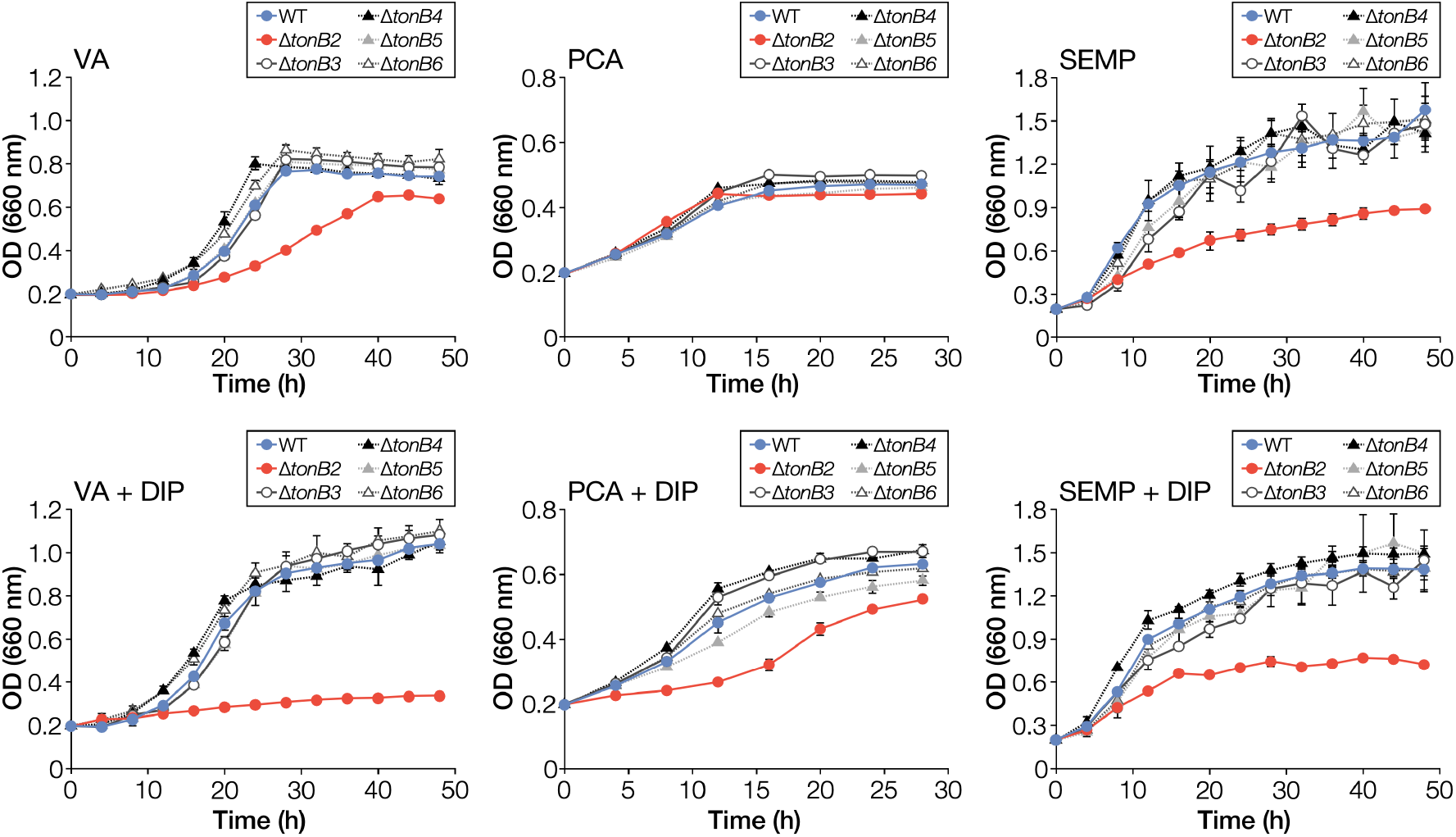
Growth of *tonB* mutants on VA, PCA, and SEMP. Cells of SYK-6, Δ*tonB2*, Δ*tonB3*, Δ*tonB4*, Δ*tonB5*, and Δ*tonB6* were cultured in Wx medium containing 5 mM VA, 5 mM PCA, or SEMP in the presence or absence of 100 μM DIP. Cell growth was monitored by measuring the OD660. Each value is the average ± the standard deviation of three independent experiments.

Δ*tonB2* retained the capacity to grow on PCA and SEMP under iron-limited conditions, implying the involvement of other *tonB* in iron acquisition. We examined the growth of Δ*tonB3456* and Δ*tonB23456* cells on VA, PCA, and SEMP under iron-limited conditions (Fig. S2). The growth of Δ*tonB3456* cells on VA and PCA was somewhat retarded compared with that of the wild type. Besides, the growth of Δ*tonB23456* cells on PCA and SEMP was lower than that of Δ*tonB2* cells indicating that any of the *tonB3-6* appears to have some involvement in iron acquisition. To evaluate the involvement of *tonB1* in iron acquisition, we introduced a plasmid carrying *tonB1* into Δ*tonB2* cells. While the growth of the *tonB2*-complemented Δ*tonB2* on VA, PCA, and SEMP under iron-limited conditions was seen to recover, the introduction of *tonB1* did not have a positive effect on the growth of Δ*tonB2* (Fig. S1). These results indicate that *tonB1* could not replace the function of *tonB2*.

We assessed the cellular localisation of TonB2 by performing western blot analysis using anti-TonB2 antibodies against a cell extract and a total membrane fraction prepared from SYK-6 grown on LB (Fig. S3). A clear signal was observed in the total membrane fraction, suggesting that TonB2 is localised in the cell membrane. The production of TonB2 in the cell membrane was also confirmed in *tonB2*-complemented Δ*tonB2* cells (Fig. S4). All these results suggest that TonB2, a component of the Ton complex, plays a vital role in growth under iron-limited conditions.

### Characterisation of a TBDR gene downstream of *tonB2*

A previous phylogenetic analysis has indicated that SLG_34550 just downstream of *tonB2* is classified into a clade comprising of known iron uptake TBDRs^15^. This gene, designated as *fiuA*, was deleted to obtain a *fiuA* mutant (Δ*fiuA*), and the capacity of the mutant to grow on VA, PCA, and SEMP was measured (Fig. S5). Δ*fiuA* cells showed growth retardation on VA and SEMP under iron-replete conditions, similar to Δ*tonB2* cells (Fig. 2) which increased further under iron-limited conditions Δ*fiuA* not growing at all on VA and growth retardation also seen on PCA. The growth of Δ*fiuA* cells on VA, PCA, and SEMP under iron-limited conditions was recovered by the introduction of a *fiuA*-carrying plasmid (Fig. S6).

**Figure 2.**
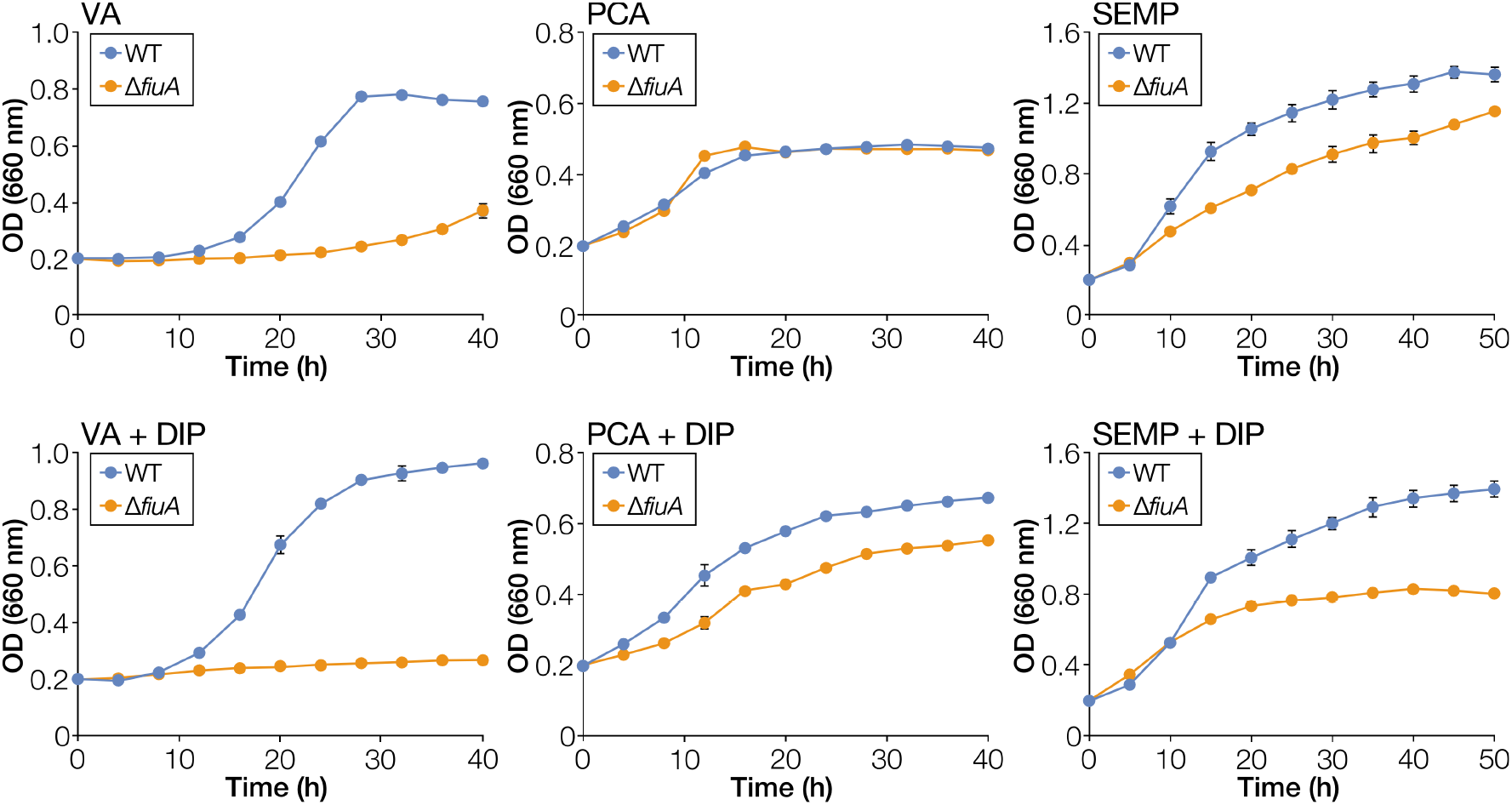
Growth of a *fiuA* mutant on VA, PCA, and SEMP. Cells of SYK-6 and Δ*fiuA* were cultured in Wx medium containing 5 mM VA, 5 mM PCA, or SEMP in the presence or absence of 100 μM DIP. Cell growth was monitored by measuring the OD660. Each value is the average ± the standard deviation of three independent experiments.

We measured the intracellular iron concentrations of Δ*tonB2*, Δ*fiuA*, and wild-type cells grown in SEMP to evaluate whether a reduction in the intracellular iron concentration resulted in the decrease in the growth of Δ*tonB2* and Δ*fiuA* (Fig. 3). The intracellular iron concentrations of Δ*tonB2* and Δ*fiuA* cells reduced to approximately 37% and 61% of that of the wild type, respectively. These results indicate the involvement of *tonB2* and *fiuA* in iron acquisition.

**Figure 3.**
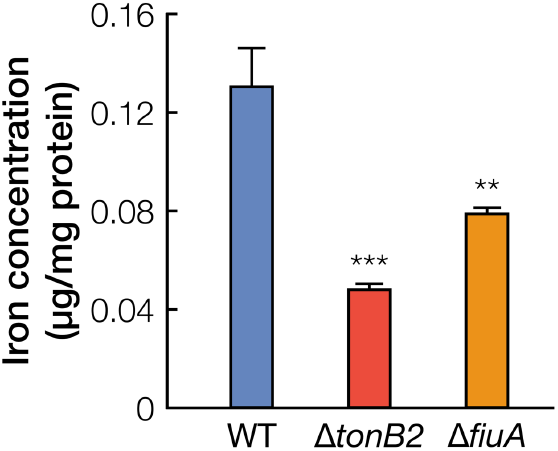
Intracellular iron concentrations of Δ*tonB2* and Δ *fiuA*. Intracellular iron concentrations of wild type, Δ*tonB2*, and Δ*fiuA* were determined as described in the Methods. Each value is the average ± the standard deviation of three independent experiments. ^**^, *P* < 0.01, ^***^, *P* < 0.001 (one-way ANOVA with Dunnett’s multiple comparisons).

### Promoter activities of *tonB2* and *fiuA* under iron-limited conditions

Rodionov et al. discovered a 19-bp Fur box sequence conserved among alphaproteobacteria using comparative genomic analysis^44^. Incomplete inverted repeat sequences similar to this Fur box sequence were found upstream of each of *tonB2* and *fiuA* (Fig. 4A, Table S1). We evaluated promoter activities of SYK-6 cells harbouring the reporter plasmid carrying a transcriptional fusion of a Fur box-containing promoter region of *tonB2* or *fiuA* with *lacZ* (Fig. 4B-D). Promoter activities were detected in both strains under iron-replete conditions, and the activities of the cells carrying the *tonB2* and *fiuA* promoter regions were seen to increase 1.6-fold and 8.0-fold, respectively under iron-limited conditions. The addition of Fe^2+^ reduced the activities to levels comparable with those of the cells under iron-replete conditions, indicating that the expression of *tonB2* and *fiuA* was induced under iron-limited conditions. However, the promoter activity of *tonB2* was significantly higher (61-fold) than that of the *fiuA* promoter under iron-replete conditions. These findings suggest that *tonB2* is expressed at a high level, even under iron-replete conditions. Western blot analysis using anti-TonB2 antibodies against the total membrane fractions obtained from SYK-6 grown with and without DIP demonstrated the production of almost equal amount of TonB2 between both membrane samples, suggesting that expression of *tonB2* is not greatly influenced by iron-limitation (Fig. S7). In contrast, the promoter activity of *tonB1* did not demonstrate any change regardless of the presence or absence of DIP (Fig. S8).

**Figure 4.**
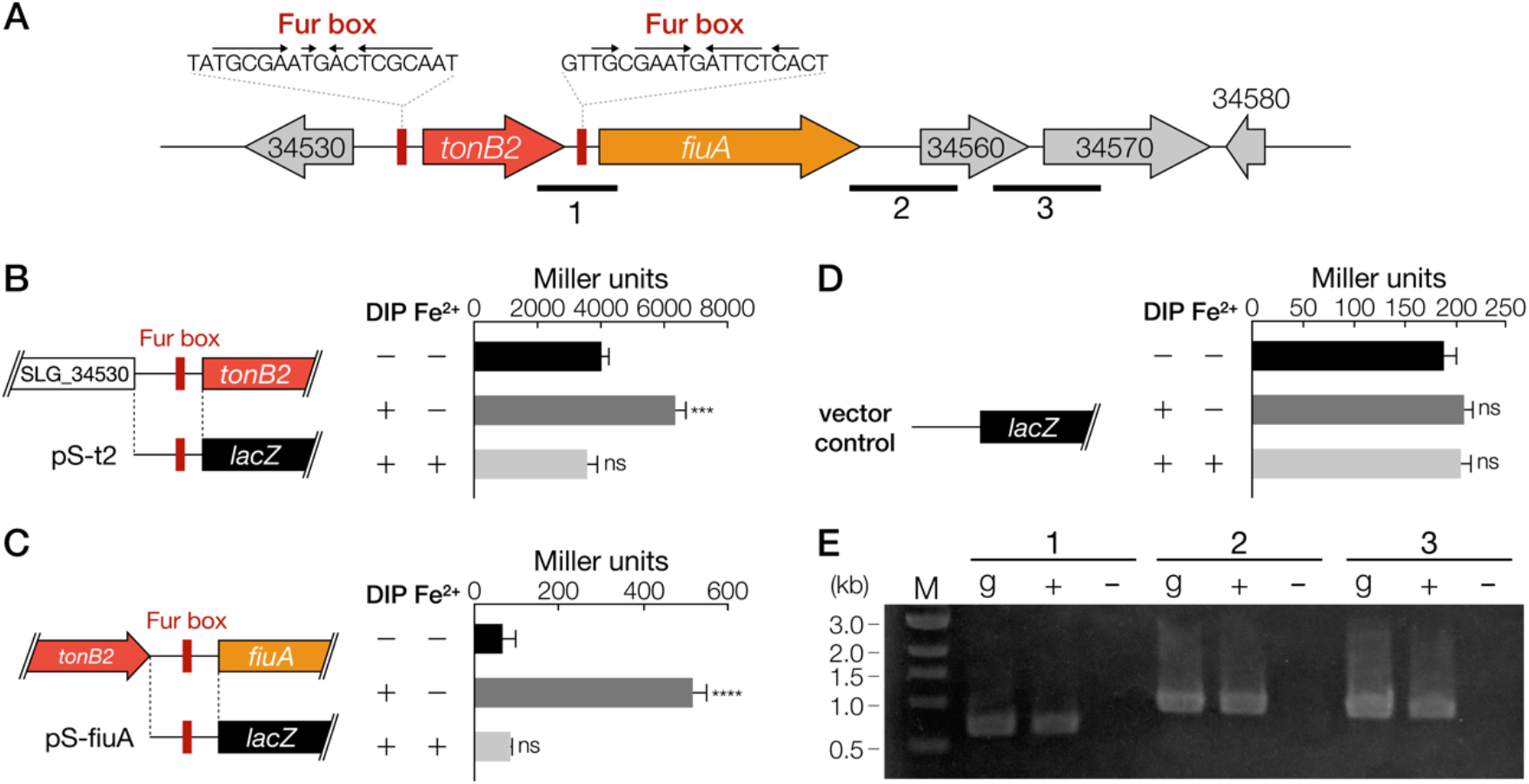
Promoter activities of *tonB2* and *fiuA* under iron-replete or limited conditions. (**A**) Gene organisation of *tonB2* and *fiuA*. Fur box-like sequences upstream of *tonB2* and *fiuA* are indicated by red squares. Genes: SLG_34530, hypothetical protein; *tonB2*, TonB-like protein; *fiuA*, TBDR; SLG_34560, putative hydroxylase; SLG_34570, putative oxidoreductase; SLG_34580, putative oxidoreductase. (**B-D**) β-galactosidase activities of SYK-6 cells harbouring pS-t2 (B), pS-fiuA (C), and pSEVA225 (D) grown in Wx-SEMP with or without 100 μM DIP and 100 μM FeCl2. The DNA fragments used for the promoter analysis are shown on the left. Each value is the average ± the standard deviation of three independent experiments. ns, *P* > 0.05, ^***^, *P* < 0.001, ^****^, *P* < 0.0001 (one-way ANOVA with Dunnett’s multiple comparisons). (**E**) RT-PCR analysis of *tonB2* and *fiuA*. Total RNA used for cDNA synthesis was isolated from SYK-6 cells grown in Wx-SEMP with 100 μM DIP. The regions to be amplified are indicated by black bars below the genetic map. Lanes: M, molecular size markers; g, control PCR with the SYK-6 genomic DNA; ‘+’ and ‘-’, RT-PCR with and without reverse transcriptase, respectively.

Although an independent promoter region was found upstream of each of *tonB2* and *fiuA*, both genes were likely to be transcribed in the same transcription unit under iron-limited conditions (Fig. 4). Reverse transcription (RT)-PCR analysis of *tonB2* and *fiuA* was performed using cDNA obtained from total RNA isolated from SYK-6 cells grown under iron-limited conditions. An amplification product between *tonB2* and *fiuA* was observed (Fig. 4E), suggesting that *fiuA* is mainly transcribed from the more active *tonB2* promoter under iron-limited conditions. These results led to the conclusion that *tonB2* and *fiuA* play significant roles in iron acquisition.

### TBDR genes other than *fiuA* required for normal growth under iron-limited conditions

Based on similarity with known iron uptake TBDRs, we found four putative iron uptake TBDRs (SLG_04340, SLG_04380, SLG_10860, and SLG_17010) in addition to *fiuA* (Table S2). These TBDR genes were classified into two phylogenetic clades, containing siderophore and haem uptake TBDRs^15^. A Fur box-like sequence was found just upstream of each gene except SLG_04340 (Table S1). RT-PCR analysis revealed that SLG_04320-SLG_04360 constituted an operon, and a Fur box-like sequence was found just upstream of SLG_04320 (Fig. S9). We evaluated promoter activities of SYK-6 cells harbouring a reporter plasmid carrying a transcriptional fusion of a Fur box-containing promoter region of SLG_04340, SLG_04380, SLG_10860, or SLG_17010 with *lacZ* (Fig. S10A-D). Promoter activities were detected in all the cells except the one harbouring a plasmid carrying the upstream region of SLG_17010. Promoter activities of the cells carrying the SLG_04340 and SLG_04380 promoter regions were seen to increase 1.6-fold and 12-fold, respectively, under iron-limited conditions (Fig. S10A, B). Next, mutants of SLG_04340, SLG_04380, and SLG_10860 were constructed and their growth was compared with the wild type on VA, PCA, and SEMP under iron-limited and replete conditions (Fig. S5, S11). Since disruption of these genes did not show significant effect on the growth of SYK-6, we constructed Δ*fiuA 4340*, Δ*fiuA 4380*, and Δ*fiuA 10860* and evaluated their growth under iron-limited conditions (Fig. S12). The growth of these double mutants on SEMP was further retarded as compared to that of the Δ*fiuA*. The growth of the former two mutants was also delayed on PCA. Thus, not only *fiuA* but also SLG_04340, SLG_04380, and SLG_10860 are involved in iron acquisition. Further, we constructed Δ*fiuA 4340 4380* and Δ*fiuA 4340 4380 10860* and evaluated their capacity to grow under iron-limited conditions (Fig. S13). When grown on SEMP, Δ*fiuA 4340 4380* cells exhibited almost the same level of growth as that of Δ*fiuA 4380*, however, Δ*fiuA 4340 4380 10860* cells showed substantial growth retardation as compared to Δ*fiuA 4340 4380*. By contrast, multiple mutations did not affect the capacity of these cells to grow on PCA, implying that different TBDRs are involved in the iron uptake during growth on PCA.

### Identification of an inner membrane iron transporter

The SYK-6 genome consists of four genes (SLG_06990, SLG_13630, SLG_36840, and SLG_p-00340) showing similarity with known inner membrane transporter genes involved in the uptake of siderophore and ferrous iron (Table S2). To evaluate their involvement in iron acquisition, we constructed mutants of these genes and measured their growth on VA, PCA, and SEMP under iron-limited and replete conditions (Fig. 5A). SLG_36840 has 28-29% amino acid sequence identity with the ferrous iron inner membrane transporter gene (*feoB*) of *Escherichia coli* K-12 (AAC76434) and *Pseudomonas aeruginosa* PAO1 (AAG07746). Δ*36840* (Δ*feoB*) showed growth retardation on VA and PCA under iron-limited conditions. The introduction of a *feoB*-carrying plasmid into Δ*feoB* recovered the growth of Δ*feoB* cells on VA and PCA under iron-limited conditions (Fig. S14). In addition, the intracellular iron concentration of Δ*feoB* cells grown on SEMP was reduced to approximately 52% of that of the wild type (Fig. 5B).

**Figure 5.**
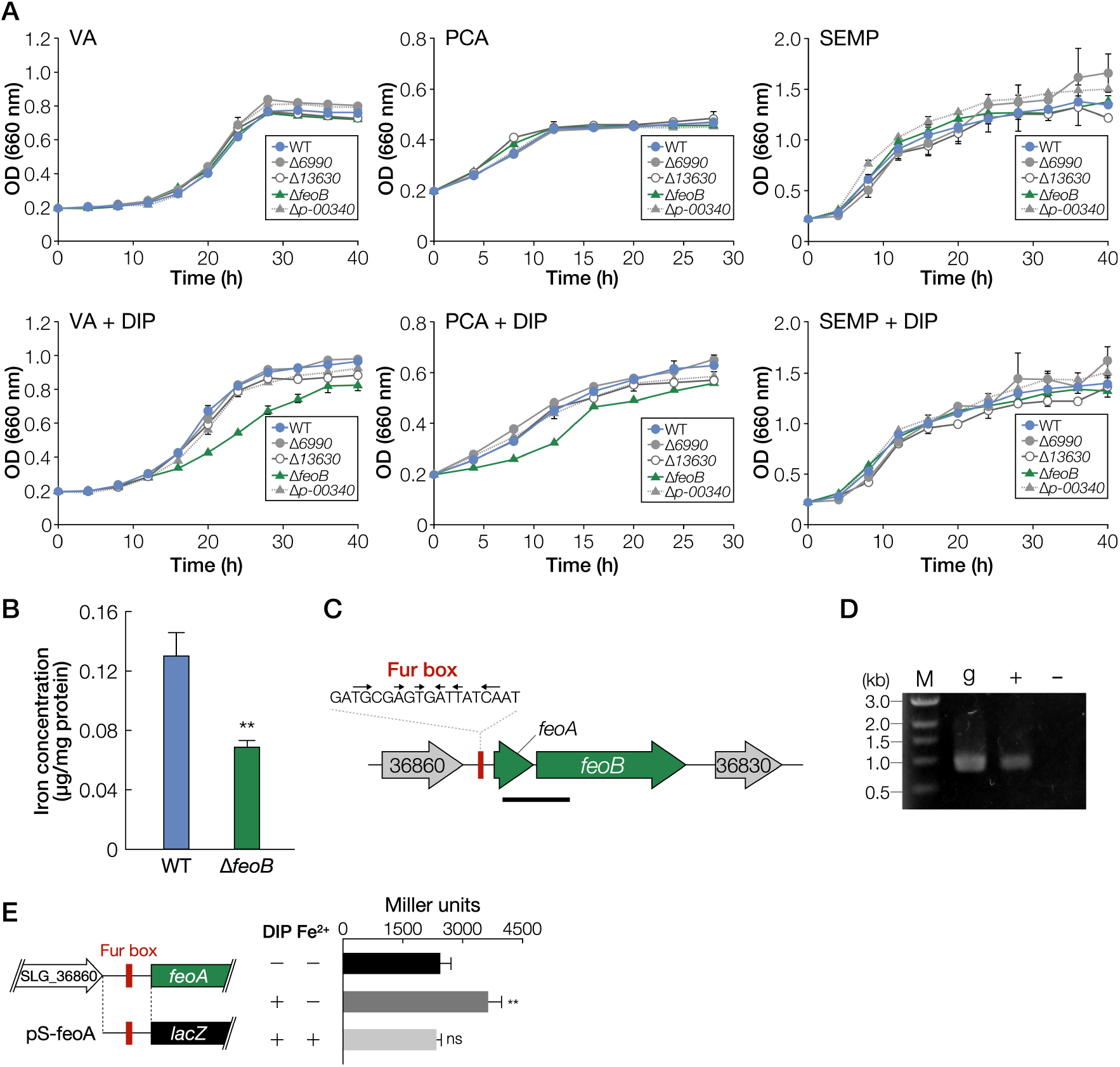
Identification of a transporter gene involved in the iron uptake across the inner membrane. (**A**) Growth of mutants of putative inner membrane iron transporter genes on VA, PCA, and SEMP. Cells of SYK-6, Δ*6990*, Δ*13630*, Δ*feoB*, and Δ*p-00340* were cultured in Wx medium containing 5 mM VA, 5 mM PCA, or SEMP in the presence or absence of 100 μM DIP. Cell growth was monitored by measuring the OD660. (**B**) Intracellular iron concentrations of wild type and Δ*feoB*. **, *P* < 0.01 (two-tailed, unpaired *t*-test). (**C**) Gene organisation of *feoAB*. Genes: SLG_36830, putative single-stranded DNA-binding protein; *feoB*, putative ferrous iron transporter protein B; *feoA*, putative ferrous iron transporter protein A; SLG_36860, putative ubiquinone biosynthesis protein. (**D**) RT-PCR analysis of *feoAB*. Total RNA used for cDNA synthesis was isolated from SYK-6 cells grown in Wx-SEMP with 100 μM DIP. The region to be amplified is indicated by a bar below the genetic map (C). Lanes: M, molecular size markers; g, control PCR with the SYK-6 genomic DNA; ‘+’ and ‘-’, RT-PCR with and without reverse transcriptase, respectively. (**E**) β-galactosidase activities of SYK-6 cells harbouring pS-feoA grown in Wx-SEMP with or without 100 μM DIP and 100 μM FeCl2. The DNA fragment used for the promoter analysis is shown on the left. Each value is the average ± the standard deviation of three independent experiments. ns, *P* > 0.05, ^**^, *P* < 0.01 (one-way ANOVA with Dunnett’s multiple comparisons).

There is a *feoA-*like gene (SLG_36850), just upstream of *feoB* that encodes an important factor for FeoB multimer formation^20^. RT-PCR analysis showed that *feoA* and *feoB* comprise an operon (Fig. 5C, D). Since a Fur box was found upstream of *feoA*, the promoter activities of a *feoA* promoter region containing the Fur box were evaluated (Fig. 5E). Promoter activity was observed to be increased 1.5-fold under iron-limited conditions. All these results indicate that *feoAB* is involved in the uptake of ferrous iron across the inner membrane. However, the growth of SYK-6 on SEMP remaining unaffected by the disruption of *feoB* is still unclear.

### Identification of Fur involved in the regulation of iron uptake genes

SYK-6 has two *fur*-like genes, *fur1* (SLG_29410) and *fur2* (SLG_05570), which showed 21% amino acid sequence identity with each other and exhibited 35% and 21%, 38% and 21%, and 68% and 20% identity with Fur of *P. aeruginosa* PAO1 (AAG08150), *E. coli* K-12 (AAC73777), and *Caulobacter crescentus* NA1000 (ACL93522), respectively. The involvement of *fur1* and *fur2* in the transcriptional regulation of *tonB2*, *fiuA*, SLG_04340, SLG_04380, and *feoAB* was evaluated by attempting to disrupt these genes, which resulted in only a *fur2* mutant being obtained (Fig. S5). We assessed the promoter activities of Δ*fur2* harbouring a reporter plasmid carrying a transcriptional fusion of each Fur box-containing promoter region of the above genes with *lacZ* (Fig. S15). However, Δ*fur2* cells harbouring each plasmid showed almost the same level of promoter activities with wild type under iron-replete conditions, indicating that *fur2* is not involved in their transcriptional regulation. Next, we examined whether Fur1 could bind these promoter regions using purified Fur1 obtained from *fur1*-expressing *E. coli* BL21(DE3) (Fig. S16). Electrophoretic mobility shift assay revealed that Fur1 was bound to the promoter regions of *tonB2*, *fiuA*, SLG_04340, SLG_04380, and *feoA* and was not bound to the promoter regions without the Fur box (Fig. 6A-E). These results strongly suggest that Fur1 regulates the expression of these genes by binding to the Fur box.

**Figure 6.**
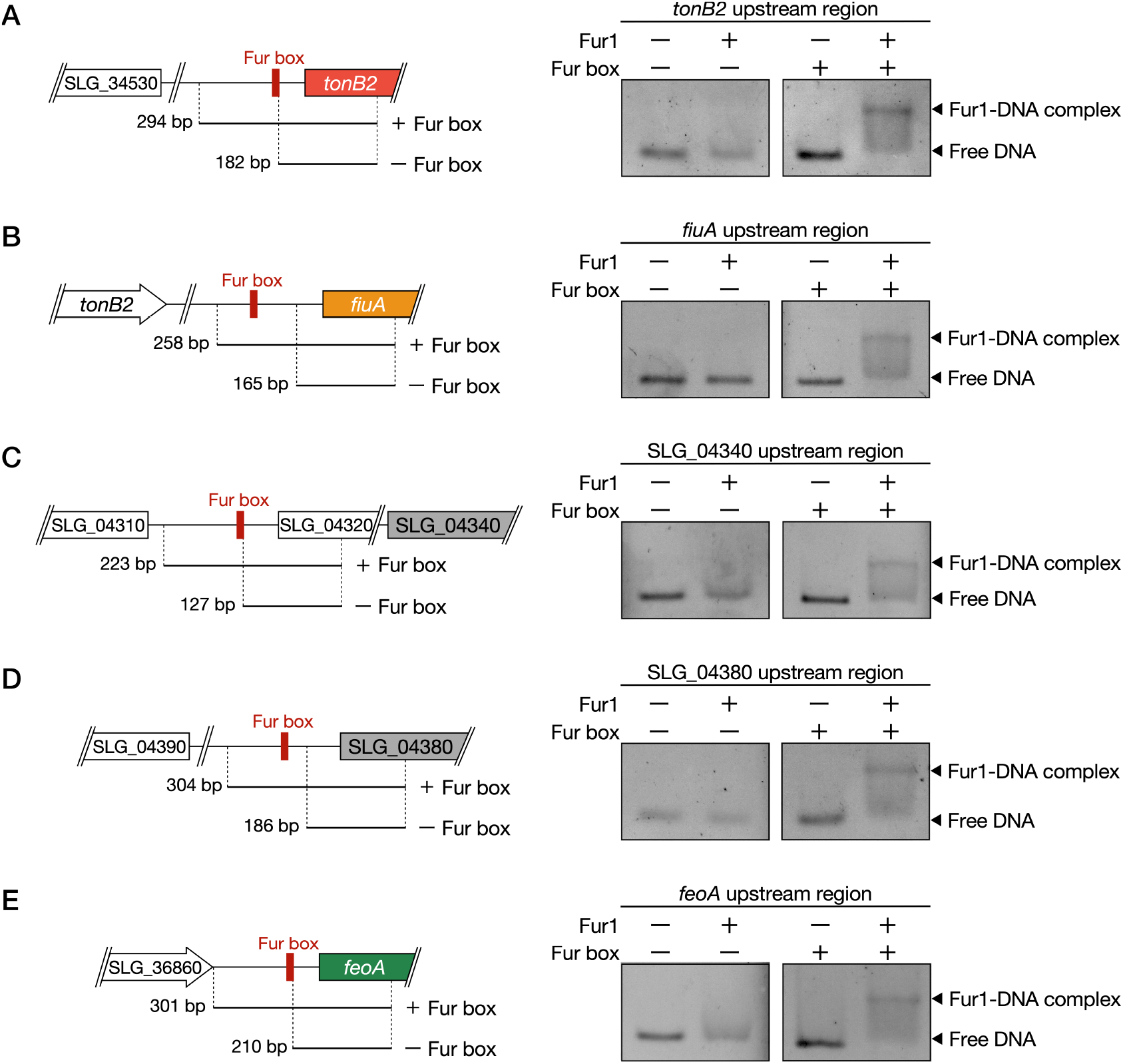
Fur1 binds to the Fur box sequences upstream of the iron uptake genes. EMSA of purified Fur1 using DNA probes of promoter regions of *tonB2* (A), *fiuA* (B), SLG_04340 (C), SLG_04380 (D), and *feoA* (E). The DNA fragments used for EMSA are shown on the left. The uncropped gel images are shown in Fig. S21.

The results of this study demonstrated that the transcription of *fiuA* was driven from the *tonB2* promoter under iron-limited conditions (Fig. 4E), and Fur1 was able to bind to the Fur box upstream of *fiuA* (Fig. 6B). Based on these results, it was hypothesised that the transcriptional regulation of *fiuA* was as described below. Under iron-replete conditions, Fur1 binds to the Fur box upstream of *fiuA*, interrupting the transcription from the *tonB2* promoter that is active even under iron-replete conditions. Under iron-limited conditions, Fur1 is released from the Fur box, and then *fiuA* is strongly co-transcribed with *tonB2*. This hypothesis was verified by constructing a plasmid carrying a transcriptional fusion of a *tonB2* promoter region and *fiuA* promoter region with *lacZ* (pS-t2-fiuA) and evaluating the promoter activities of SYK-6 cells harbouring pS-t2-fiuA (Fig. S17A). Under iron-limited conditions, the promoter activities between the cells harbouring pS-t2-fiuA and the cells harbouring pS-t2, which carries a *tonB2* promoter-*lacZ* fusion, were almost the same. Under iron-replete conditions, while the activity of SYK-6 cells harbouring pS-t2 decreased to only ca. 63% of the activity under iron-limited conditions, the activity of SYK-6 cells harbouring pS-t2-fiuA was drastically reduced (ca. 11%). These observations, thus, support the hypothesis (Fig. S17B). Based on the *tonB2* expression profile (highly expressed regardless of both iron-replete and -limited conditions), TonB2 likely interacts with not only FiuA but also other TBDRs to acquire iron and other nutrients.

### PCA is a potential siderophore used for SYK-6

SYK-6 cells were grown on Wx-SEMP agar medium containing chrome azurol S (CAS) to examine whether SYK-6 secretes siderophores (Fig. S18). SYK-6 cells grown for 144 h formed a slight halo around the colony, suggesting that SYK-6 cells weakly secrete siderophores. However, no genes showed similarity with known siderophore synthetase genes in the SYK-6 genome. Besides, the size of the halo did not change when Δ*tonB2* and Δ*fiuA* were assayed. These results suggest that SYK-6 does not promote producing and secreting siderophores even under iron-limited conditions, unlike other siderophore-producing bacteria^11^.

As shown in Fig. 1 and Fig. 2, Δ*tonB2* and Δ*fiuA* did not exhibit severe growth retardation on PCA as compared to VA and SEMP. This fact may imply that the TonB2-FiuA-independent iron acquisition system functions in SYK-6 during its growth on PCA. Since PCA is known to form a complex with iron^45,46^, we examined whether the addition of PCA improves the growth of Δ*tonB2* and Δ*fiuA* on SEMP under iron-limited conditions (Fig. 7, Fig. S19). Interestingly, the addition of 100 μM PCA improved their growth after 40 h as seen from the OD_660_ values of the cultures which was 1.3- to 1.4-fold higher than that without PCA with high concentrations of PCA (500 to 1000 μM) showing more effect on promoting growth (1.6- to 1.8-fold). We measured the growth of a *fiuA ligAB* double mutant (Δ*fiuA ligAB*) to confirm that PCA did not contribute to these growth improvements as a carbon source. *ligAB* encodes PCA 4,5-dioxygenase which is essential for growth of SYK-6 on PCA^47^. Δ*fiuA ligAB* cells exhibited increased growth (1.8- to 2.5-fold) when PCA (100 to 1000 μM) was added (Fig. 7). These results suggest that SYK-6 cells utilize PCA as a siderophore or secrete unknown siderophore(s) induced by PCA. Since Δ*tonB2* and Δ*fiuA* showed reduced growth on VA which is metabolised via PCA^47^ and the presence of PCA did not promote halo formation in CAS assay, PCA appears to act as a siderophore (Fig. S18). Notably, the growth improvement of SYK-6 and Δ*ligAB* cells on SEMP was modest with the addition of PCA, unlike Δ*fiuA ligAB* cells (Fig. 7). These results suggest that TonB2 and FiuA are mainly involved in the iron acquisition and that the pathway utilizing PCA as a siderophore is ancillary.

**Figure 7.**
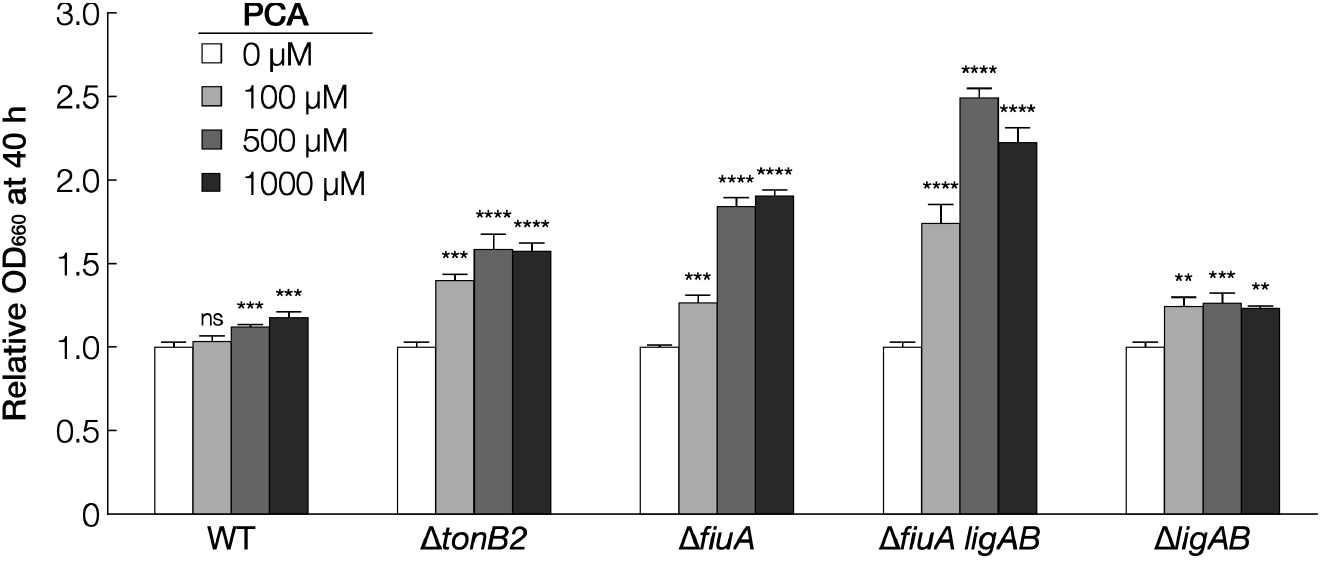
PCA enhances the growth of Δ *tonB2* and Δ *fiuA* under iron-limited conditions. Cells of SYK-6, Δ*tonB2*, Δ*fiuA*, Δ*fiuA ligAB*, and Δ*ligAB* were cultured in Wx-SEMP containing 100 μM DIP with or without PCA (100 μM, 500 μM, or 1000 μM). Cell growth was monitored by measuring the OD660. Each bar shows a relative value of OD660 at 40 h of cultures in the presence of PCA when the OD660 at 40 h of culture in the absence of PCA was set to 1.0 (leftmost bars). Each value is the average ± the standard deviation of three independent experiments. ns, *P* > 0.05, ^**^, *P* < 0.01, ^***^, *P* < 0.001, ^****^, *P* < 0.0001 (one-way ANOVA with Dunnett’s multiple comparisons). Individual growth curves are shown in Fig. S19.

### Iron uptake genes of SYK-6 are partially conserved in Sphingomonadaceae

We investigated whether the genes involved in iron uptake in SYK-6 are conserved in ten Sphingomonad strains shown in Table S3. It has been reported that approximately 40% of Gram-negative bacteria with known genomes have more than two *tonB*-like genes^48^. The Sphingomonad strains compared here have 3-8 *tonB*-like genes, however, there was no gene showing >40% amino acid sequence identity with *tonB2*. The proportion of Gram-negative bacteria, which have more than 30 TBDR genes in their genomes, is only ca. 16%^14^. Since the Sphingomonad strains have a large number of TBDR genes (from 39 to 153), they may play important roles not only in iron acquisition but also in other functions^15,49^. Five of the ten strains investigated showed the presence of TBDR-like genes, which showed 49-54% amino acid sequence identity with *fiuA*. Although their similarities with *fiuA* were somewhat low, a phylogenetic tree of all TBDRs of the five strains demonstrated the formation of a specific clade containing *fiuA* and its homologs mentioned above with known iron uptake TBDRs (Fig. S20). Among them, there was a *tonB* homolog in the vicinity of the *fiuA*-like gene of *Novosphingobium nitrogenifigens* DSM 19370, *Novosphingobium* sp. PP1Y, and *N. pentaromativorans* US6-1, respectively. Thus, these *fiuA* and *tonB* homologs are likely to participate in the outer membrane iron uptake. In contrast, every Sphingomonad strain contained genes whose amino acid sequence showed 32-60% and 62-72% identities with those of SYK-6 *feoA* and *feoB*. FeoAB may therefore play an important role in the inner membrane ferrous iron uptake in Sphingomonadaceae.

This study demonstrated that the Ton system plays a major role in iron uptake across the outer membrane in SYK-6 and that the Feo system is necessary for its normal growth. SYK-6 appears to uptake ferric iron across the outer membrane using the Ton system consisting of *tonB2* and *fiuA*. Ferric iron is probably reduced to ferrous iron in the periplasm, and then it is taken up by the Feo system. In addition to the Feo system, ferric iron appears to be incorporated into the cytoplasm using unidentified transporters (e.g. ABC transporters). The reduction of ferric iron in the periplasm is essential for ferrous iron uptake by the Feo system, however, genes similar to periplasmic reductase such as *vciB* of *V. cholerae* were not found in the SYK-6 genome^50,51^. Thus, it would be very important to identify the periplasmic ferric reductase and alternative inner membrane iron transporter genes and to clarify whether FiuA takes up free iron or ferric complex.

## Methods

### Bacterial strains, plasmids, culture conditions, and substrates

The strains and plasmids (Table S4) and the PCR primers (Table S5) were used in this study. *Sphingobium* sp. SYK-6 (NBRC 103272/JCM 17495) and its mutants were grown at 30 °C with shaking (160 rpm) in LB or Wx minimal medium containing SEMP^43^. Media for SYK-6 transformants and mutants was supplemented with 50 mg l^−1^ kanamycin (Km). *E*. *coli* strains were cultured in LB at 37 °C. Media for *E*. *coli* transformants carrying antibiotic resistance markers was supplemented with 25 mg l^−1^ Km or 100 mg l^−1^ ampicillin (Amp).

### Mutant construction

Plasmids for gene disruption were constructed by amplifying ca. 1-kb fragments carrying upstream and downstream regions of each gene by PCR with SYK-6 genome DNA as a template and the primer pairs as shown in Table S5. The resultant fragments were inserted into the BamHI site in pAK405 by In-Fusion cloning (TaKaRa Bio, Inc.). These plasmids were independently introduced into SYK-6 cells and its mutants by triparental mating, and candidate mutants were isolated as previously described^52^. Disruption of the genes was confirmed by colony PCR using primer pairs (Table S5). The plasmids for gene complementation of Δ*tonB2*, Δ*fiuA*, and Δ*feoB* (Table S4) were introduced into the mutants by electroporation.

### Sequence analysis

Sequence analysis was performed using the MacVector program version 15.5.2. Sequence similarity searches, multiple alignments, and pairwise alignments were performed using the BLAST program^53^, Clustal Omega program^54^, and the EMBOSS program^55^, respectively. A phylogenetic tree was generated using the FigTree program (http://tree.bio.ed.ac.uk/software/figtree/).

### RT-PCR analysis

SYK-6 cells grown in LB were harvested and washed twice with Wx medium. The cells were resuspended to an optical density at 600 nm (OD_600_) of 0.2 in Wx-SEMP and cultured at 30 °C until OD_600_ of the culture reached 0.5, after which they were incubated in the presence of 100 μM DIP for 2 h. Total RNA was isolated from the cells using an Illumina RNAspin Mini RNA isolation kit (GE Healthcare). The samples were treated with DNase I to remove any contaminating genomic DNA. Total RNA (4 μg) was reverse transcribed using SuperScript IV reverse transcriptase with random hexamer primers.

The cDNA was purified using a NucleoSpin Gel and PCR Clean-up kit (Takara Bio, Inc.). PCR was performed with the cDNA, specific primers (Table S5), and Gflex DNA polymerase (Takara Bio, Inc.). The DNA obtained was electrophoresed on a 0.8% agarose gel.

### Growth measurement

SYK-6 cells, its mutants, and complemented strains were grown in LB for 24 h. The cells were harvested by centrifugation at 4,800 × *g* for 5 min, washed twice with Wx medium, and resuspended in 3 ml of the same medium. The cells were then inoculated in Wx medium containing SEMP, 5 mM VA, or 5 mM PCA to an OD_660_ of 0.2 with or without 100 μM DIP. Since SYK-6 exhibits auxotrophy for methionine when grown in a methoxy-group-free substrate, 20 mg l^−1^ methionine was added to the medium to grow on PCA. Cells were incubated at 30 °C with shaking (60 rpm) and cell growth was monitored every hour by measuring the OD_660_ with a TVS062CA biophotorecorder (Advantec Co., Ltd.).

The complemented strains of Δ*feoB*, Δ*fiuA*, and Δ*tonB2* were analysed by growing cells in Wx medium containing Km and 1 mM *m*-toluate (an inducer of the P*m* promoter in pJB861).

### Promoter assay

SYK-6 cells and Δ*fur2* harbouring each plasmid (Table S4) grown in LB containing Km for 20 h were harvested by centrifugation at 4,800 × *g* for 5 min, washed twice with Wx medium, and resuspended in 1 ml of the same medium. The cells were then inoculated in Wx-SEMP containing Km to an OD_600_ of 0.2. Samples were incubated at 30 °C with shaking (60 rpm) until OD_600_ of the culture reached 0.5. Then, the cells were further incubated with or without 100 μM DIP and 100 μM FeCl_2_ for 2 h. β-galactosidase activity of the cells was measured using 2-nitrophenyl-β-D-galactopyranoside as described previously^15^ and expressed as Miller units.

### Western blot analysis

A peptide corresponding to residues 247–266 (HGPDPRDRPLSDGQIKTIET) of TonB2 was synthesised and used as an antigen to obtain antisera against TonB2 in rabbits (Cosmo Bio, Inc.). Anti-TonB2-peptide antibodies were obtained by purification of the antiserum using peptide affinity column chromatography (Cosmo Bio, Inc.). Total membrane fractions were prepared as described previously from SYK-6 cells incubated in LB for 20 h with or without 100 μM DIP^15^. When total membrane fractions were prepared from the *tonB2*-complemented Δ*tonB2*, cells were incubated in LB containing Km and 1 mM *m*-toluate. TonB2 was detected by western blot analysis using anti-TonB2 antibodies (0.09 μg/ml) as described previously^15^. Horseradish peroxidase-conjugated goat anti-rabbit IgG antibodies (Invitrogen, 0.2 μg/ml) were used as the secondary antibodies. Protein concentrations were determined by the Bradford method using a Bio-Rad protein assay kit or Lowry’s assay with a DC protein assay kit (Bio-Rad Laboratories). TonB2 was detected using the ECL Western Blotting Detection System (GE Healthcare) with a LumiVision PRO image analyser (Aisin Seiki Co., Ltd).

### Intracellular iron concentration measurement

SYK-6 cells and its mutants were grown in LB for 20 h, harvested by centrifugation at 4,800 × *g* for 5 min, washed twice with Wx medium, and resuspended in 1 ml of the same medium. The cells were then inoculated in Wx medium containing SEMP to an OD_600_ of 0.2. Samples were incubated at 30 °C with shaking (60 rpm) for 6 h. The cells were harvested by centrifugation, washed twice with 50 mM Tris-HCl buffer (pH 7.5), and resuspended in 200 μl of the same buffer. The cells were disrupted by sonication to obtain cell lysates. The cell lysates were then centrifuged at 18,800 × *g* for 10 min and the protein concentration of the resulting supernatants (cell extracts) was determined. The iron concentration of cell extracts was determined using an Iron Assay Kit LS (Metallogenics Co., Ltd.) based on the ferrozine chromogenic method. Protein concentrations were determined using a Bio-Rad protein assay kit.

### Expression of *fur1* in *E. coli* and purification of Fur1

A *fur1*-coding region was PCR-amplified from SYK-6 genome DNA using primers listed in Table S5. A 0.4-kb NdeI-BamHI fragment carrying *fur1* was inserted into the corresponding sites of pET-16b (pET-fur1) by In-Fusion cloning (TaKaRa Bio, Inc.). *E. coli* harbouring pET-fur1 was grown in LB containing Amp at 30 °C until the OD_600_ of the culture reached 0.5, and then the expression of *fur1* was induced for 4 h at 30 °C by addition of 1 mM isopropyl-β-D-thiogalactopyranoside. The cells were harvested by centrifugation at 4,800 × *g* for 5 min, washed twice with 50 mM Tris-HCl buffer (pH 7.5) containing 100 mM NaCl, and resuspended in 200 μl of the same buffer. The cells were disrupted by sonication and cell lysate was obtained. The cell lysate was then centrifuged at 18,800 × *g* for 10 min and the resulting supernatant was applied to His Spin Trap (GE Healthcare). After centrifugation (100 × *g*, 1 min, 4 °C), samples were washed 3 times with 50 mM Tris-HCl buffer (pH 7.5) containing 100 mM NaCl and 50 mM imidazole, and Fur1 was eluted with 50 mM Tris-HCl buffer (pH 7.5) containing 100 mM NaCl and 500 mM imidazole. Purified Fur1 was subjected to desalting and concentrating by centrifugal filtration using an Amicon Ultra 3k (Millipore). The purity of Fur1 was analysed by sodium dodecyl sulfate-15% polyacrylamide gel electrophoresis. Protein concentrations were determined by a Bio-Rad protein assay kit.

### Electrophoretic mobility shift assay

DNA probes were PCR-amplified from SYK-6 genome DNA using the primers listed in Table S5. The DNA-protein binding reactions were performed at 20 °C for 30 min in 10 μl of binding buffer (50 mM Tris-HCl, 5 mM dithiothreitol, 50 mM MgCl_2_, 200 mM KCl, and 0.5 % [wt/vol] Tween 20, pH 7.5) containing 20 fmol DNA probe, 500 ng of purified Fur1, 1 μg of poly(dI-dC), and 100 mM MnSO_4_. The resulting samples were separated by electrophoresis on 2.5% agarose gel and signals were detected using SYBR™ Gold Nucleic Gel Stain (Invitrogen).

### CAS assay

Fifty millilitres of 1.2 g l^-1^ chrome azurol S solution was mixed with 10 ml of 1 mM FeCl_3_ (dissolved in 10 mM HCl) and 40 ml of 5 mM hexadecyltrimethylammonium bromide (CAS solution). Eighteen millilitres of a Wx-SEMP agar medium with or without PCA (final conc. 1 mM) was mixed with 2 ml of the CAS solution to prepare CAS assay plates. The cells of SYK-6, Δ*tonB2*, and Δ*fiuA* were grown in LB for 20 h, harvested by centrifugation at 4,800 × *g* for 5 min, washed twice with Wx medium, and resuspended in 1 ml of the same medium. Ten microlitres of the culture (OD_600_ = 10) was inoculated on a cellulose filter (12 mm) on a CAS assay plate and incubated for 6 days at 30 °C.

### Statistics and Reproducibility

All results were obtained from *n* = 3 independent experiments. Statistical tests were performed using Graphpad Prism8 (Graphpad software). One-way ANOVA with Dunnett’s multiple comparisons and unpaired, two-tailed *t*-test were used as shown in figure legends. *P* < 0.05 was considered statistically significant.

## Data availability

All data supporting this study are available within the article and its Supplementary Information or are available from the corresponding author upon request.

## Acknowledgements

We thank Yudai Higuchi and Aya Takeuchi for assistance with the construction of the SLG_06990, SLG_13630, and SLG_p-00340 mutants. This work was supported by JSPS KAKENHI Grant Numbers 15H04473, 19H02867, and 19J11312.

## Author Contributions

E.M. supervised the project. M.F., N.K., and E.M. designed the study and wrote the manuscript. M.F. performed data analysis, western blot analysis, intracellular iron concentration measurement, purification of Fur1, EMSA, and CAS assay. M.F. and T.S. constructed plasmids and performed growth measurement and promoter assay. T.S., M.F., H.Y. and K.M. constructed mutants. K.T. performed RT-PCR analysis and helped to perform EMSA. T.S., K.T., H.Y., and K.M. helped to interpret the data and discussed the results. All authors read and approved the manuscript.

