## Supplemental files for "Iron acquisition system of *Sphingobium* sp. strain SYK-6, a degrader of lignin-derived aromatic compounds"

**Table S1. Fur box-like sequences upstream of *tonB2*, TBDR genes, and *feo* in SYK-6**

| Locus tag | Putative function | Accession number | Fur box-like sequence | Position (from start codon) |
|---|---|---|---|---|
| SLG_04320–04360 | TBDR | BAK65109.1 | GCTGCAACTCACTATCAAT | −37 to −19 |
| SLG_04380 | TBDR | BAK65113.1 | AATGCGAATCATTCGCATT | −121 to −103 |
| SLG_10860 | TBDR | BAK65761.1 | AATGATATTAACTCGCAAT | −61 to −43 |
| SLG_17010 | TBDR | BAK66376.1 | GTTGCGACTAACTCGCATT | −43 to −25 |
| SLG_34540 (*tonB2*) | TonB | BAK68129.1 | TATGCGAATGACTCGCAAT | −47 to −29 |
| SLG_34550 (*fiuA*) | TBDR | BAL68130.1 | GTTGCGAATGATTCTCACT | −89 to −71 |
| SLG_36840 (*feoB*) | FeoB | BAK68359.1 | ATTGATAATCACTCGCATC | −39 to −21 |

### Table S2. SYK-6 genes showing homology with known iron transporters

| Protein | Function[a] | Substrate | Accession no. | Strain | Similar genes in SYK-6 | Sequence identity (%)[b] | Putative function | Refer-ence |
|---|---|---|---|---|---|---|---|---|
| **Inner membrane** | | | | | | | | |
| FepB | SBP | Enterobactin | P0AEL6 | *Escherichia coli* K12 | – | – | – | 10 |
| FhuD | SBP | Ferrichrome | P07822 | *Escherichia coli* K12 | – | – | – | 11 |
| FecB | SBP | Ferric citrate | P15028 | *Escherichia coli* K12 | – | – | – | 12 |
| SirA | SBP | Staphyloferrin B | BAB41330 | *Staphylococcus aureus* N315 | – | – | – | 13 |
| FatB | SBP | Ferric anguibactin | P11460 | *Vibrio anguillarum* 775 | – | – | – | 14 |
| PhuT | SBP | Haem | AAC13287 | *Pseudomonas aeruginosa* PAO1 | – | – | – | 15 |
| ShuT | SBP | Haem | O70018 | *Shigella dysenteriae* | – | – | – | 16 |
| FptX | Transporter | Pyochelin | AAG07606 | *Pseudomonas aeruginosa* PAO1 | SLG_13630 | 22 | AmpG | 17 |
| RhtX | Transporter | Rhizobactin 1021 | Q92XI7 | *Sinorhizobium meliloti* 2011 | – | – | – | 17 |
| FeoB | Transporter | $Fe^{2+}$ | P33650 | *Escherichia coli* K-12 | SLG_36840 | 29 | FeoB | 18 |
| | | | AAG07746 | *Pseudomonas aeruginosa* PAO1 | SLG_36840 | 28 | FeoB | 19 |
| | | | | | SLG_p-00340 | 11 | High-affinity iron transporter | |
| EfeU | Transporter | $Fe^{2}$ | P75901 | *Escherichia coli* K-12 | – | – | – | 20 |
| MntH | Transporter | $Mn^{2+}$, $Fe^{2+}$ | P0A769 | *Escherichia coli* K-12 | – | – | – | 21 |
| ZupT | Transporter | $Zn^{2+}$, $Co^{2+}$, $Fe^{2+}$ | P0A8H3 | *Escherichia coli* K-12 | SLG_06990 | 31 | ZIP family transporter | 22 |
| FutA1 | SBP | $Fe^{2+}$ | P72827 | *Synechocystis* sp. strain PCC 6803 | – | – | – | 23 |
| FutA2 | SBP | $Fe^{2+}$ | Q55835 | *Synechocystis* sp. strain PCC 6803 | – | – | – | 24 |
| YfeA | SBP | $Fe^{2+}$ | Q56952 | *Yersinia pestis* | – | – | – | 25 |
| **Outer membrane** | | | | | | | | |
| FhuA | TBDR | Ferrichrome | P06971 | *Escherichia coli* K-12 | SLG_17010 | 26 | TBDR | 26 |
| FhuE | TBDR | Coprogen | P16869 | *Escherichia coli* K-12 | SLG_04340 | 22 | TBDR | 27 |
| Fiu | TBDR | Catecholate siderophore | P75780 | *Escherichia coli* K-12 | SLG_34550 | 30 | TBDR | 28 |
| FoxA | TBDR | Ferrioxamine | Q9I116 | *Pseudomonas aeruginosa* PAO1 | SLG_17010 | 23 | TBDR | 29 |
| FpvA | TBDR | Pyoverdine | P48632 | *Pseudomonas aeruginosa* PAO1 | SLG_10860 | 21 | TBDR | 30 |
| FptA | TBDR | Pyochelin | P42512 | *Pseudomonas aeruginosa* PAO1 | SLG_17010 | 27 | TBDR | 31 |
| PupA | TBDR | Pseudobactin 358 | P25184 | *Pseudomonas putida* WCS358 | SLG_34550 | 20 | TBDR | 32 |
| FcuA | TBDR | Ferrichrome | Q05202 | *Yersinia enterocolitica* | SLG_04340 | 34 | TBDR | 33 |
| TbpA | TBDR | Transferrin | AHW76545 | *Neisseria meningitidis* | SLG_04380 | 22 | TBDR | 34 |
| ShuA | TBDR | Haem | P72412 | *Shigella dysenteriae* | SLG_04380 | 25 | TBDR | 35 |
| HutA | TBDR | Haem | ACL95742 | *Caulobacter crescentus* NA1000 | SLG_04380 | 47 | TBDR | 36 |
| HasR | TBDR | Haemophore | Q79AD2 | *Serratia marcescens* | SLG_04380 | 21 | TBDR | 37 |

[a]SBP, Substrate binding protein of ABC transporter; TBDR, TonB-dependent receptor.
[b]Amino acid sequence identity was calculated using EMBOSS Needle pairwise alignment program.

**Table S3. Genes most similar to SYK-6 *tonB2*, *fiuA*, and *feoAB* in Sphingomonads**

| Strain | Number of *tonB* | Number of TBDR | TonB2 | | FiuA | | FeoA | | FeoB | |
|---|---|---|---|---|---|---|---|---|---|---|
| | | | Accession no. | Sequence identity (%)[a] | Accession no. | Sequence identity (%)[a] | Accession no. | Sequence identity (%)[a] | Accession no. | Sequence identity (%)[a] |
| *Blastomonas natatoria* | 3 | 55 | PXW77659 | 20 | PXW77661 | 49 | PXW76439 | 34 | PXW76438 | 69 |
| *Novosphingobium aromaticivorans* DSM12444 | 3 | 76 | ABD24466 | 26 | ABD27715 | 28 | ABD27286 | 41 | ABD27285 | 63 |
| *Novosphingobium nitrogenifigens* DSM 19370 | 4 | 76 | EGD58708 | 34 | EGD58710 | 28 | EGD57881 | 38 | EGD57882 | 62 |
| *Novosphingobium pentaromativorans* US6-1 | 4 | 68 | EHJ58335 | 35 | EHJ60384 | 32 | EHJ60269 | 34 | EHJ60270 | 62 |
| *Novosphingobium* sp. PP1Y | 3 | 98 | CCA92020 | 22 | CCA92021 | 33 | CCA92126 | 37 | CCA92127 | 62 |
| *Sphingobium chlorophenolicum* L-1 | 5 | 94 | AEG50735 | 33 | AEG50736 | 49 | AEG49960 | 60 | AEG49961 | 70 |
| *Sphingobium japonicum* UT26S | 4 | 69 | BAI95867 | 38 | BAI95866 | 49 | BAI95131 | 54 | BAI95132 | 72 |
| *Sphingobium yanoikuyae* ATCC 51230 | 8 | 100 | EKU76051 | 35 | EKU72862 | 54 | EKU74251 | 48 | EKU74252 | 70 |
| *Sphingomonas wittichii* RW1 | 3 | 153 | ABQ70391 | 35 | ABQ69571 | 51 | ABQ68809 | 35 | ABQ68810 | 68 |
| *Sphingopyxis alaskensis* RB2256 | 3 | 39 | ABF54651 | 27 | ABF53800 | 26 | ABF54691 | 32 | ABF54692 | 68 |

[a]Amino acid sequence identity was calculated using EMBOSS Needle pairwise alignment program.

### Table S4. Strains and plasmids used in this study

| Strains or plasmids | Relevant characteristic(s)[a] | Reference or source |
|---|---|---|
| Strains | | |
| *Sphingobium* sp. | | |
| SYK-6 | Wild type; Nal$^r$ Sm$^r$ | 1 |
| SME096 | SYK-6 derivative; ΔSLG_36940 (*tonB3*) | 2 |
| SME097 | SYK-6 derivative; ΔSLG_34540 (*tonB2*) | 2 |
| SME151 | SYK-6 derivative; ΔSLG_13630 | This study |
| SME219 | SYK-6 derivative; ΔSLG_04340 | This study |
| SME220 | SYK-6 derivative; ΔSLG_04380 | This study |
| SME221 | SYK-6 derivative; ΔSLG_10860 | This study |
| SME225 | SYK-6 derivative; ΔSLG_34550 (*fiuA*) | This study |
| SME257 | SYK-6 derivative; ΔSLG_12500 12510 (*ligAB*) | 3 |
| SME290 | SYK-6 derivative; ΔSLG_37490 (*tonB4*) | 2 |
| SME292 | SYK-6 derivative; ΔSLG_01650 (*tonB5*) | 2 |
| SME293 | SYK-6 derivative; ΔSLG_14690 (*tonB6*) | 2 |
| SME303 | SYK-6 derivative; Δ*tonB3456* | 2 |
| SME304 | SYK-6 derivative; Δ*tonB23456* | 2 |
| SME305 | SYK-6 derivative; ΔSLG_04340 SLG_04380 *fiuA* | This study |
| SME306 | SYK-6 derivative; ΔSLG_04340 *fiuA* | This study |
| SME307 | SYK-6 derivative; ΔSLG_04380 *fiuA* | This study |
| SME309 | SYK-6 derivative; ΔSLG_05570 (*fur2*) | This study |
| SME310 | SYK-6 derivative; ΔSLG_36840 (*feoB*) | This study |
| SME311 | SYK-6 derivative; ΔSLG_10860 *fiuA* | This study |
| SME312 | SYK-6 derivative; Δ*ligAB fiuA* | This study |
| SME313 | SYK-6 derivative; ΔSLG_04340 SLG_04380 SLG_10860 *fiuA* | This study |
| SME314 | SYK-6 derivative; ΔSLG_p-00340 | This study |
| SME315 | SYK-6 derivative; ΔSLG_06990 | This study |
| *Escherichia coli* | | |
| HB101 | *recA13 supE44 hsd20 ara-14 proA2 lacY1 galK2 rpsL20 xyl-5 mtl-1* | 4 |
| NEB 10-beta | *araD*139 Δ(*ara-leu*)7697 *fhuA lacX74 galK* (ϕ80 Δ*lacZ* Δ*M15*) *recA1 endA1 nupG rpsL* (Sm$^r$) Δ(*mrr-hsdRMS-mcrBC*) | New England Biolabs |
| BL21 (DE3) | F$^-$ *ompT hsdSB*($r_B^-$ $m_B^-$) *gal dcm* (DE3); T7 RNA polymerase gene under control of the *lacUV5* promoter | 5 |
| Plasmids | | |
| pRK2013 | Tra$^+$ Mob$^+$ ColE1 replicon; Km$^r$ | 6 |
| pJB861 | RK2 ori broad-host-range expression vector; Km$^r$ $P_m$ *xylS* | 7 |
| pAK405 | Plasmid for allelic exchange and markerless gene deletions in Sphingomonads; Km$^r$ | 8 |
| pSEVA225 | RK2 *ori lacZ* promoter probe broad host range vector; Km$^r$ | 9 |
| pET-16b | Expression vector; T7 promoter, Ap$^r$ | Novagen |
| pJB-fiuA | pJB861 with a 2.3-kb BamHI fragment carrying *fiuA* | This study |
| pS-tonB2 | pSEVA338 with a 0.9-kb fragment carrying *tonB2* | This study |
| pJB-tonB2 | pJB861 with a 0.9-kb NotI-SacI fragment carrying *tonB2* from pS-tonB2 | This study |
| pJB-tonB1 | pJB861 with a 0.7-kb NotI-SacI fragment carrying *tonB1* | 2 |
| pJB-feoB | pJB861 with a 1.9 -kb BamHI fragment carrying *feoB* | This study |
| pAK4320 | pAK405 with a 1.9-kb deletion cassette carrying up- and downstream regions of SLG_04320 | This study |
| pAK4380 | pAK405 with a 2.0-kb deletion cassette carrying up- and downstream regions of SLG_04380 | This study |
| pAKfur2 | pAK405 with a 2.2-kb deletion cassette carrying up- and downstream regions of *fur2* | This study |
| pAK6990 | pAK405 with a 2.2-kb deletion cassette carrying up- and downstream regions of SLG_06990 | This study |
| pAK10860 | pAK405 with a 2.0-kb deletion cassette carrying up- and downstream regions of SLG_10860 | This study |
| pAK13630 | pAK405 with a 2.2-kb deletion cassette carrying up- and downstream regions of SLG_13630 | This study |

| | | |
|---|---|---|
| pAKfur1 | pAK405 with a 2.3-kb deletion cassette carrying up- and downstream regions of SLG_29410 (*fur1*) | This study |
| pAKfiuA | pAK405 with a 2.0-kb deletion cassette carrying up- and downstream regions of *fiuA* | This study |
| pAKfeoB | pAK405 with a 2.2-kb deletion cassette carrying up- and downstream regions of *feoB* | This study |
| pAKp-00340 | pAK405 with a 2.2-kb deletion cassette carrying up- and downstream regions of SLG_p-00340 | This study |
| pS-4340 | pSEVA225 with a 0.2-kb PCR amplicon carrying SLG_04340 promoter region | This study |
| pS-4380 | pSEVA225 with a 0.4-kb PCR amplicon carrying SLG_04380 promoter region | This study |
| pS-10860 | pSEVA225 with a 0.3-kb PCR amplicon carrying SLG_10860 promoter region | This study |
| pS-17010 | pSEVA225 with a 0.2-kb PCR amplicon carrying SLG_17010 promoter region | This study |
| pS-t2 | pSEVA225 with a 0.4-kb PCR amplicon carrying *tonB2* promoter region | This study |
| pS-fiuA | pSEVA225 with a 0.2-kb PCR amplicon carrying *fiuA* promoter region | This study |
| pS-t2-fiuA | pS-t2 with a 0.2-kb HindIII fragment carrying *fiuA* promoter region | This study |
| pS-feoA | pSEVA225 with a 0.1-kb PCR amplicon carrying *feoA* promoter region | This study |
| pS-t1 | pSEVA225 with a 0.2-kb PCR amplicon carrying *tonB1* promoter region | This study |
| pET-fur1 | pET-16b with a 0.4-kb NdeI-BamHI fragment carrying *fur1* | This study |

[a]Nal$^r$, Sm$^r$, Km$^r$, and Amp$^r$, resistance to nalidixic acid, streptomycin, kanamycin, and ampicillin, respectively.

### Table S5. Primers used in this study

| Target gene | Primer | Sequences (5′ to 3′) |
|---|---|---|
| For gene disruption | | |
| pAK4340 | Dis_TopF | CGGTACCCGGGGATCGATGGGCTTGCTGTTCCTC |
| | Dis_TopR | GAAGCAGGGTCGGGAAAC |
| | Dis_BotF | GTTTCCCGACCCTGCTTCCCTTCGATGCGTTCAACCA |
| | Dis_BotR | CGACTCTAGAGGATCCTCCAGCTCCTCGGCATAA |
| pAK4380 | Dis_TopF | CGGTACCCGGGGATCACATCATCGCCCAGCTCG |
| | Dis_TopR | TATGCGTGGTGGAGCGAC |
| | Dis_BotF | GTCGCTCCACCACGCATAAGAGCCCGTCATTGTCCT |
| | Dis_BotR | CGACTCTAGAGGATCGCCAAGAGCATCTGCAAGA |
| pAK5570 | Dis_TopF | CGGTACCCGGGGATGTCCTCCGGCGTGTTGAA |
| (*fur2*) | Dis_TopR | GAACGCCCCGTGATCGAA |
| | Dis_BotF | TTCGATCACGGGGCGTTCGCCCTTGCCTTCAAGCGT |
| | Dis_BotR | CGACTCTAGAGGATCGTGAGGAGCCGCGTCATT |
| pAK6990 | Dis_TopF | ATTCGAGCTCGGTACCCGGGAAGCCGGGCCAAGATCGT |
| | Dis_TopR | GTCGGGGATGATGATGTTGTCCCTCGCTGG |
| | Dis_BotF | ACAACATCATCATCCCCGACATGCATGAC |
| | Dis_BotR | CCTGCAGGTCGACTCTAGAGCATCTTTTCGCCACCACG |
| pAK10860 | Dis_TopF | CGGTACCCGGGGATCGATGACTTCTGCGGCTGTA |
| | Dis_TopR | AGCGATGACCATAAACGAGT |
| | Dis_BotF | ACTCGTTTATGGTCATCGCTCATGGGCAACCTCAACAACC |
| | Dis_BotR | CGACTCTAGAGGATCCATGCTGCGGCGATGAAC |
| pAK13630 | Dis_TopF | CGGTACCCGGGGATCGAAAGCAGGCTTGTCGGTC |
| | Dis_TopR | GAGCAGGAAGGCGACGAA |
| | Dis_BotF | TTCGTCGCCTTCCTGCTCCTTCTGGGCCTTCACGGT |
| | Dis_BotR | CGACTCTAGAGGATCGTATCGGGTTCAGCCTTCG |
| pAK29410 | Dis_TopF | CGGTACCCGGGGATAGCATGCAGGCCACCGAT |
| (*fur1*) | Dis_TopR | AATCACACGGCGCTGCTC |
| | Dis_BotF | GAGCAGCGCCGTGTGATTATCGCGAGCGCAAGGACT |
| | Dis_BotR | CGACTCTAGAGGATCTGCGGACCGACGACGATA |
| pAK34550 | Dis_TopF | CGGTACCCGGGGATCCTTCATGCACCCCAGCTTC |
| (*fiuA*) | Dis_TopR | CGACACAGGAAAGGGCAAG |
| | Dis_BotF | CTTGCCCTTTCCTGTGTCGGCAGCTCAACGTGAAGAACT |
| | Dis_BotR | CGACTCTAGAGGATCTTGTCCGCCCTCATGTCT |
| pAK36840 | Dis_TopF | CGGTACCCGGGGATGTCGTTGCCAGCCGTTTCG |
| (*feoB*) | Dis_TopR | TGCCTGGTTCGTCTTCGC |
| | Dis_BotF | GCGAAGACGAACCAGGCAGGCTTGCGGCGATGAGAA |
| | Dis_BotR | CGACTCTAGAGGATCAAACGCCGCCTTCGATGG |
| pAKp-00340 | Dis_TopF | CGGTACCCGGGGATGCCAGTCTCTCCTCCACA |
| | Dis_TopR | TAGTCGAGCAGCCGCCAT |
| | Dis_BotF | ATGGCGGCTGCTCGACTAGCAACTCCTGATGGCGGT |
| | Dis_BotR | CGACTCTAGAGGATCTGGCGTCGGCATTGATAC |
| For confirmation of gene disruption | | |
| SLG_04340 | | |
| SLG_04380 | Conf_F | TGAATCCCCGATCCTGACC |
| *fur2* | Conf_F | TGTGATCGAGCGGGCTTT |
| SLG_06990 | Conf_F | GCCTGCGCGAACACATTG |
| SLG_10860 | Conf_F | CTACATGTCGCTGAACGCG |
| SLG_13630 | Conf_F | CCACTCCAGATGAAGCATGT |
| SLG_17010 | Conf_F | ATCATGCGCTCGATATCCCG |
| *fiuA* | Conf_F | TTGGCTTTCCCCCGATTG |
| *feoB* | Conf_F | GGTTGGGGTATGAACGCA |

| | | |
|---|---|---|
| SLG_p-00340 | Conf_F | TCATGCCGTCAGCACGAC |
| | Conf_F | GATCGGCTTGAGGGTGGC |
| For RT-PCR | | |
| 04320-04330 | Forward | GCGAGCTGGCGGAAATGA |
| | Reverse | GAGTCGCCCAGGAAATCG |
| 04330-04340 | Forward | CGATTTCCTGGGCGACTC |
| | Reverse | GCATCGTGGTGTAGGTCC |
| 04340-04350 | Forward | AACAAGGCGCAGGGCATT |
| | Reverse | GCGGATAGTCAACAGAGCC |
| 04350-04360 | Forward | GGCTCTGTTGACTATCCGC |
| | Reverse | CGGTCGGCACATAATGGA |
| *feoA-feoB* | Forward | ACGCATGATCGTGGGGAT |
| | Reverse | GCAACGCGGCGAAACTCT |
| *tonB2-fiuA* | Forward | GTGGACCGCAGGATATGG |
| | Reverse | CACAGGAAAGGGCAAGGA |
| *fiuA*-34560 | Forward | AGCATCACCGCCAACTAC |
| | Reverse | GGGAAGATCGCATTGTCG |
| 34560-34570 | Forward | CGACAATGCGATCTTCCC |
| | Reverse | GTCTGTTCGCGCGTTTCC |
| For EMSA | | |
| SLG_04320 | Forward (+Fur box) | TACCCGCTCGATTTGCCTC |
| | Forward (-Fur box) | ATGACGAGTGCGGGATCAG |
| | Reverse | GTAGAGCCCGTCATTGTCC |
| SLG_04380 | Forward (+Fur box) | GTGAGGGCGTATGCAGAGG |
| | Forward (-Fur box) | GCGAACACTGAAGGTCCGT |
| | Reverse | TGGAAGATCAACGGCAGGC |
| *tonB2* | Forward (+Fur box) | CGGCTCCGAATATCTCTGC |
| | Forward (-Fur box) | TCGGTTGGGGTATGAACGC |
| | Reverse | GAAGCTGGGGTGCATGAAG |
| *fiuA* | Forward (+Fur box) | CGCATCCTCTACAGATCGG |
| | Forward (-Fur box) | CAACAGCAGACAGGGGAAC |
| | Reverse | GGTATCGGTGACGGTGAC |
| *feoB* | Forward (+Fur box) | GACTGCGGGTCGATAGCC |
| | Forward (-Fur box) | AGAAAGCTCCGTCTTGCG |
| | Reverse | ATCCCCACGATCATGCGT |
| For plasmid construction | | |
| pJB-fiuA | Forward | GAAGCTTCGTGGATCTCCAGTCGAACAACCGTG |
| | Reverse | CAGGATATCTGGATCCATGAGCATTTCCTTCGCC |
| pS-tonB2 | Forward | GCCTAGGCCGCGGCCGCGCGATTACAGGACCGGGCACG |
| | Reverse | ATCCCCGGGTACCGAGCTCGTCAGGTCTCGATCGTCTTG |
| pJB-feoB | Forward | GAAGCTTCGTGGATCTCGGCCTTGCCGCGTCAG |
| | Reverse | CAGGATATCTGGATCCTAGAGGCCCAGGGCCAC |
| pS-4320 | Forward | CCTCTAGAGTCGACCGGAAACCTCCAGACCAGA |
| | Reverse | CTAAGCTTGCATGCCGGACCTTCAGTGTTCGCT |
| pS-4380 | Forward | CCTCTAGAGTCGACCGAGCGTCGCGGCGCCGAT |
| | Reverse | CTAAGCTTGCATGCCGCGCGCGGTTCCACGGTC |
| pS-10860 | Forward | CCTCTAGAGTCGACCGCTGTTAGCACTCACCCT |
| | Reverse | CTAAGCTTGCATGCCGAATAGCCCCCGAAAATGT |
| pS-17010 | Forward | CCTCTAGAGTCGACCCCTGACGATCGAGGCGGC |
| | Reverse | CTAAGCTTGCATGCCGACAGCCCCCTCGGAAGA |
| pS-t2 | Forward | CCTCTAGAGTCGACCCGGCGCCGGCGCAGAAGC |
| | Reverse | CTAAGCTTGCATGCCACCCCAACCGATCGTGCC |

| | | |
|---|---|---|
| pS-fiuA | Forward | CCTCTAGAGTCGACCGCGCGGCCGGCGCGGGCA |
| | Reverse | CTAAGCTTGCATGCCTGGTTCCCCTGTCTGCTG |
| pS-t2-fiuA | Forward | ATCGGTTGGGGTGGCATGCAGCGCGGCCGGCGCGGGCA |
| | Reverse | GTCATATGTTTTTCCTCCTATGGTTCCCCTGTCTGCTGTTGTGCATTTCGCCGACGC |
| pS-feoA | Forward | CCTCTAGAGTCGACCAAGGTCCGACTGCGGGTC |
| | Reverse | CTAAGCTTGCATGCCAAGACGGAGCTTTCTTCGA |
| pS-t1 | Forward | CCTCTAGAGTCGACCTAAGATGATTCTCCGGAGG |
| | Reverse | CTAAGCTTGCATGCCTTCAGCAACGAAACTCCTTA |
| pET-fur1 | Forward | GGCCATACGAAGGTCGTCATACCCGGACAATCGATATTGAAGC |
| | Reverse | TCGGGCTTTGTTAGCAGCCGTCAGTCCTTGCGCTCGCG |

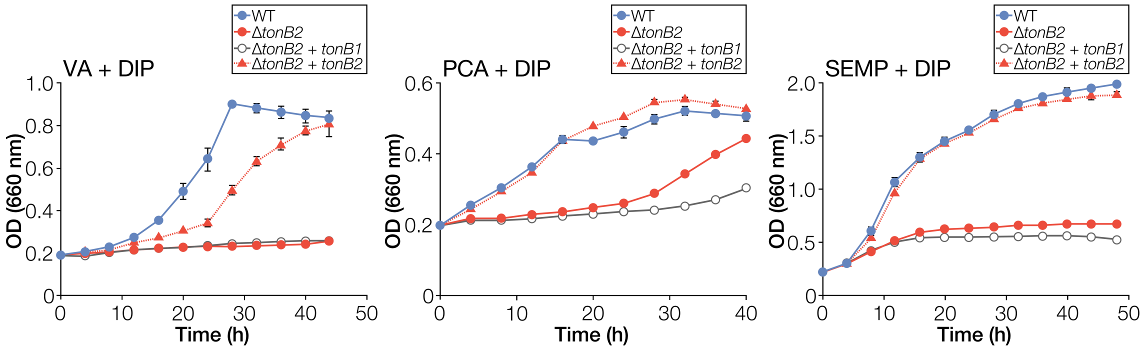


**Fig. S1. Growth complementation of Δ*tonB2* under iron-limited conditions.** Cells of SYK-6(pJB861, vector), Δ*tonB2*(pJB861), Δ*tonB2*(pJB-tonB1), and Δ*tonB2*(pJB-tonB2) were cultured in Wx medium containing 5 mM VA, 5 mM PCA, or SEMP with 100 μM DIP and 1 mM *m*-toluate. Cell growth was monitored by measuring the $OD_{660}$. Each value is the average ± the standard deviation of three independent experiments.

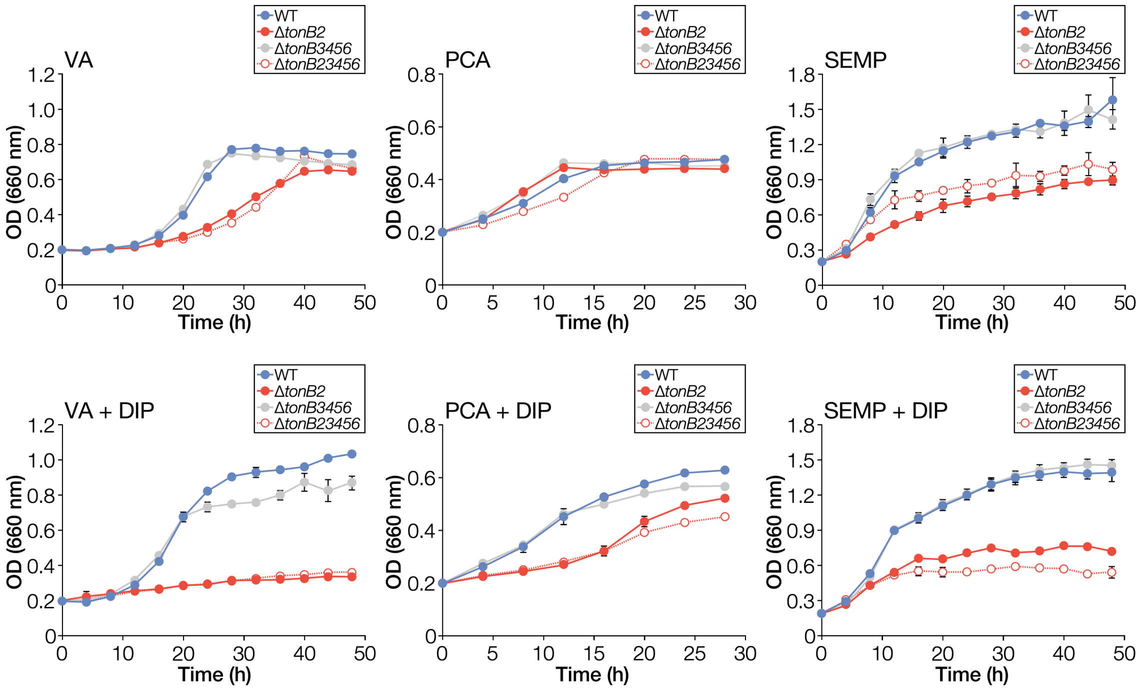


**Fig. S2. Growth of *tonB* multiple mutants on VA, PCA, and SEMP.** Cells of SYK-6, Δ*tonB2*, Δ*tonB3456*, and Δ*tonB23456* were cultured in Wx medium containing 5 mM VA, 5 mM PCA, or SEMP in the presence or absence of 100 µM DIP. Cell growth was monitored by measuring the $OD_{660}$. Each value is the average ± the standard deviation of three independent experiments.

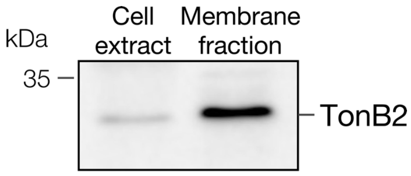


**Fig. S3. Cellular localisation of TonB2.** Western blot analysis using anti-TonB2 antibodies was performed against cell extract and total membrane fraction (10 µg protein) obtained from SYK-6 cells grown in LB. The uncropped blot image is shown in Fig. S21.

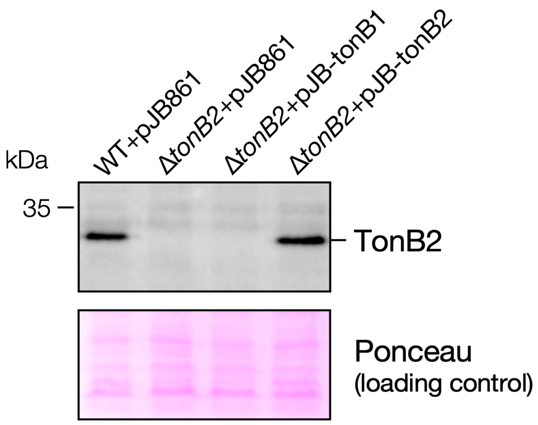


**Fig. S4. Expression of *tonB2* in a *tonB2*-complemented Δ*tonB2*.** Western blot analysis using anti-TonB2 antibodies was performed against total membrane fractions (10 µg protein) obtained from SYK-6(pJB861, vector), Δ*tonB2*(pJB861), Δ*tonB2*(pJB-tonB1), and Δ*tonB2*(pJB-tonB2) cells grown in LB containing 1 mM *m*-toluate. Ponceau staining is shown as loading control. The uncropped blot and ponceau staining image are shown in Fig. S21.

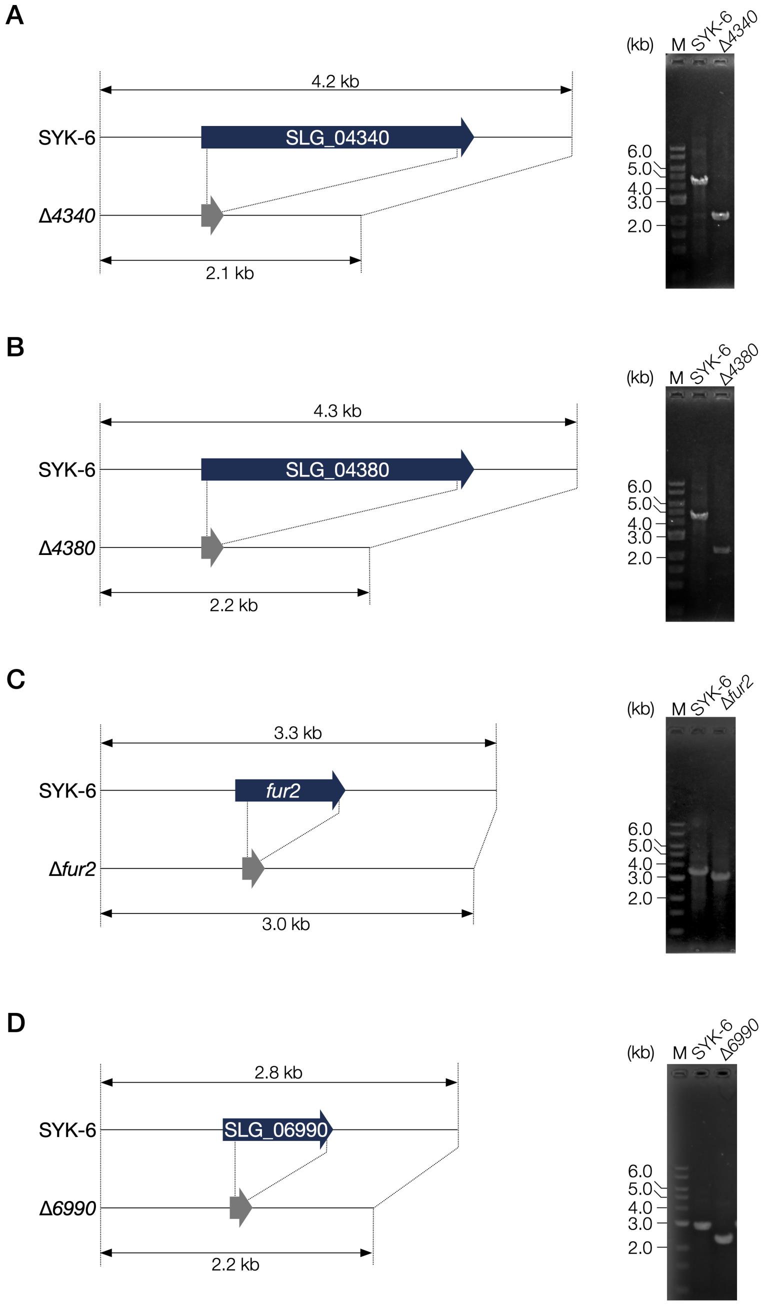


**Fig. S5. Construction of mutants.** Schematic representations and colony PCR analyses of the disruption of SLG_04340 (A), SLG_04380 (B), *fur2* (C), SLG_06990 (D), SLG_10860 (E), SLG_13630 (F), *fiuA* (G), *feoB* (H), and SLG_p-00340 (I). The primer pairs used for colony PCR analyses are shown in Table S5 (Conf_F and Dis_BotR). M, molecular size markers.

E

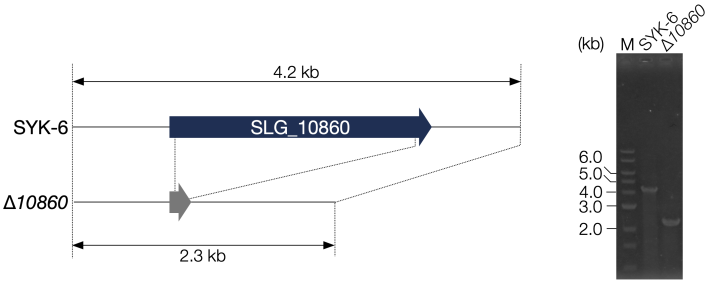


F

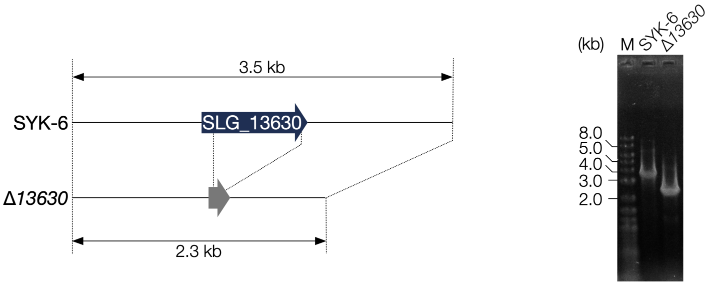


G

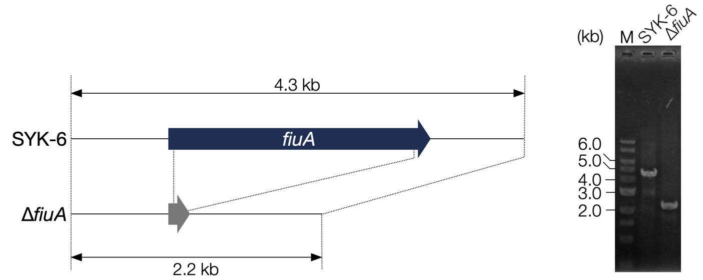


H

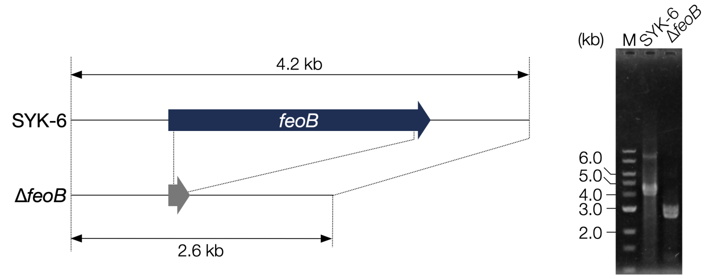


I

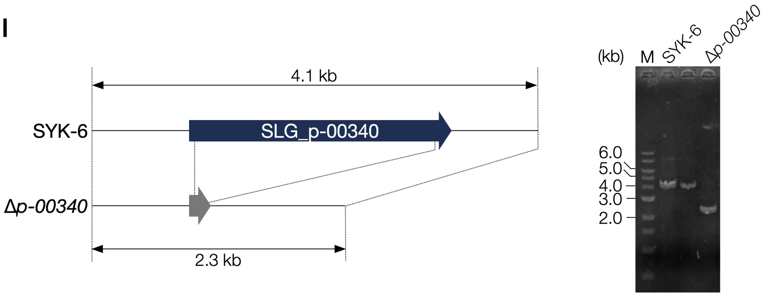


**Fig. S5.** *–continued.*

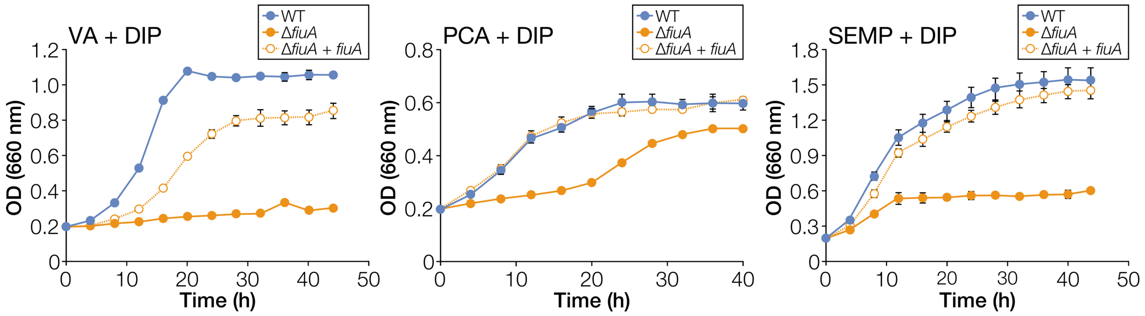


**Fig. S6. Growth complementation of Δ*fiuA* under iron-limited conditions.** Cells of SYK-6(pJB861, vector), Δ*fiuA*(pJB861), and Δ*fiuA*(pJB-fiuA) were cultured in Wx medium containing 5 mM VA, 5 mM PCA, or SEMP with 100 μM DIP and 1 mM *m*-toluate. Cell growth was monitored by measuring the $OD_{660}$. Each value is the average ± the standard deviation of three independent experiments.

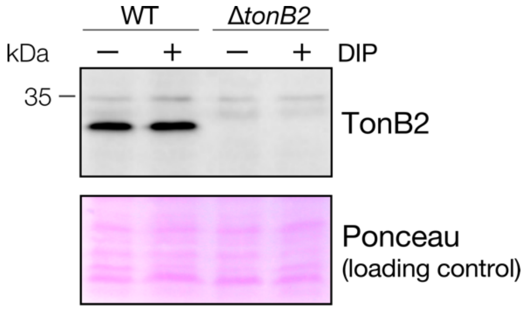


**Fig. S7. Level of TonB2 production under iron-limited conditions.** Western blot analysis using anti-TonB2 antibodies was performed against total membrane fractions (10 μg protein) obtained from SYK-6 and Δ*tonB2* cells grown in LB with or without 100 μM DIP. The band intensities of TonB2 determined by LumiVision PRO image analyser (Aisin Seiki Co., Ltd) were 222,851,628 (WT – DIP) and 258,797,925 (WT + DIP), respectively. The uncropped blot and ponceau staining image are shown in Fig. S21.

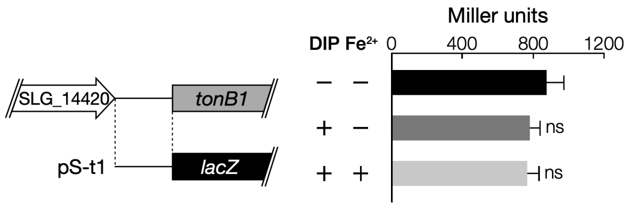


**Fig. S8. Promoter activities of *tonB1* under iron-replete and limited conditions.** β-galactosidase activities of SYK-6 cells harbouring pS-t1 grown in Wx-SEMP with or without 100 μM DIP and 100 μM $FeCl_2$ are shown. The DNA fragments used for the promoter analysis are shown on the left. Each value is the average ± the standard deviation of three independent experiments. ns, $P > 0.05$ (one-way ANOVA with Dunnett's multiple comparisons).

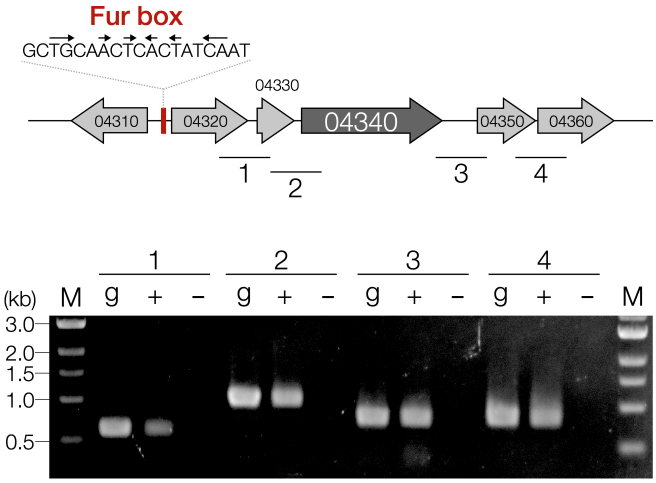


**Fig. S9. RT-PCR analysis of the SLG_04320–04360 gene cluster.** Total RNA used for cDNA synthesis was isolated from SYK-6 cells grown in Wx-SEMP with 100 μM DIP. The regions to be amplified are indicated by black bars below the genetic map. Lanes: M, molecular size markers; g, control PCR with the SYK-6 genomic DNA; '+' and '–', RT-PCR with and without reverse transcriptase, respectively.

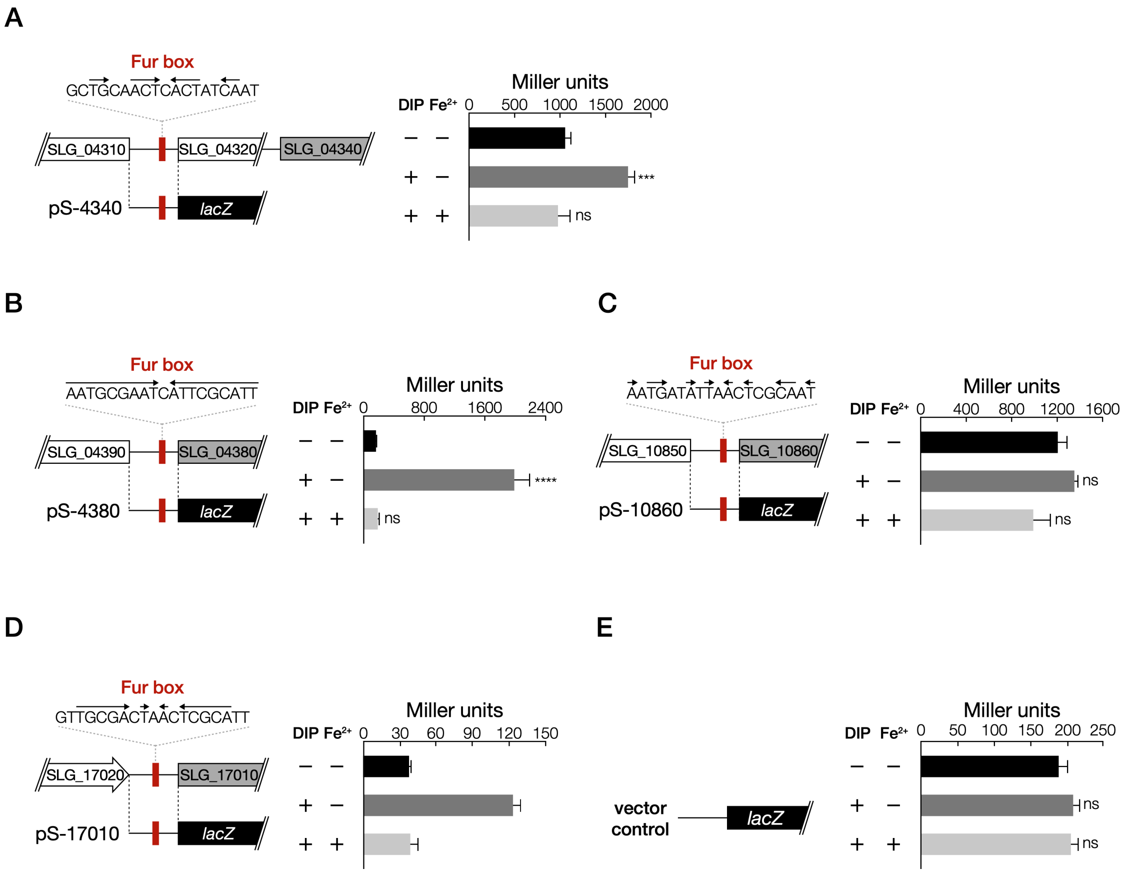


**Fig. S10. Promoter activities of TBDR candidate genes involved in iron uptake under iron-replete and limited conditions**. β-galactosidase activities of SYK-6 cells harbouring pS-4340 (A), pS-4380 (B), pS-10860 (C), pS-17010 (D), or pSEVA225 (E) grown in Wx-SEMP with or without 100 µM DIP and 100 µM $FeCl_2$. The DNA fragments used for the promoter analysis are shown on the left. Each value is the average ± the standard deviation of three independent experiments. ns, $P > 0.05$, ***, $P < 0.001$, ****, $P < 0.0001$ (one-way ANOVA with Dunnett's multiple comparisons).

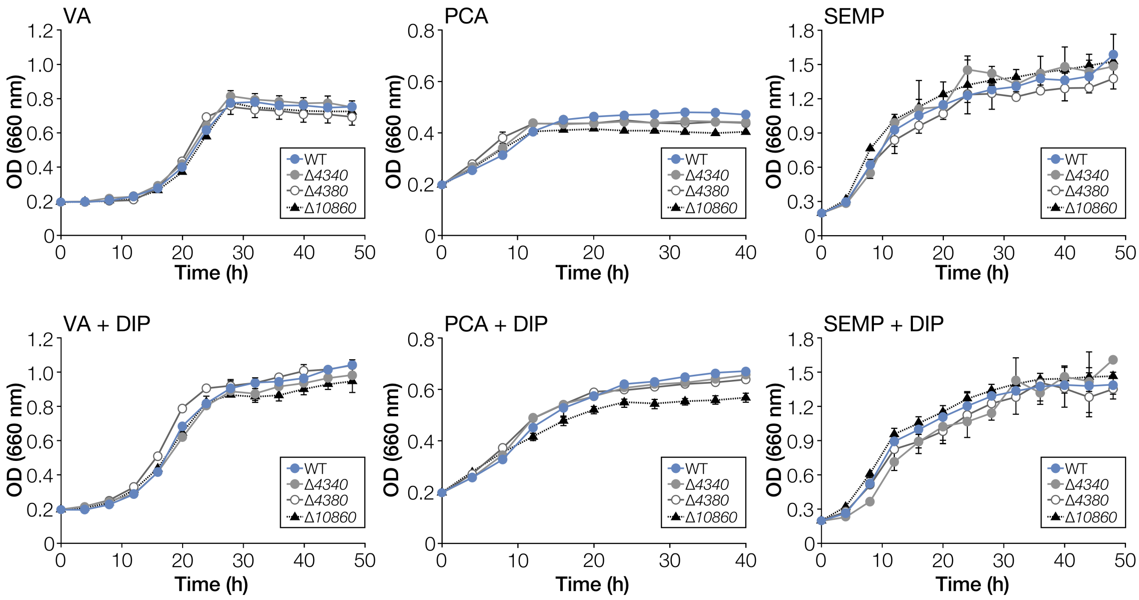


**Fig. S11. Growth of Δ*4340*, Δ*4380*, and Δ*10860* on VA, PCA, and SEMP.** Cells of SYK-6, Δ*4340*, Δ*4380*, and Δ*10860* were cultured in Wx medium containing 5 mM VA, 5 mM PCA, or SEMP in the presence or absence of 100 µM DIP. Cell growth was monitored by measuring the $OD_{660}$. Each value is the average ± the standard deviation of three independent experiments.

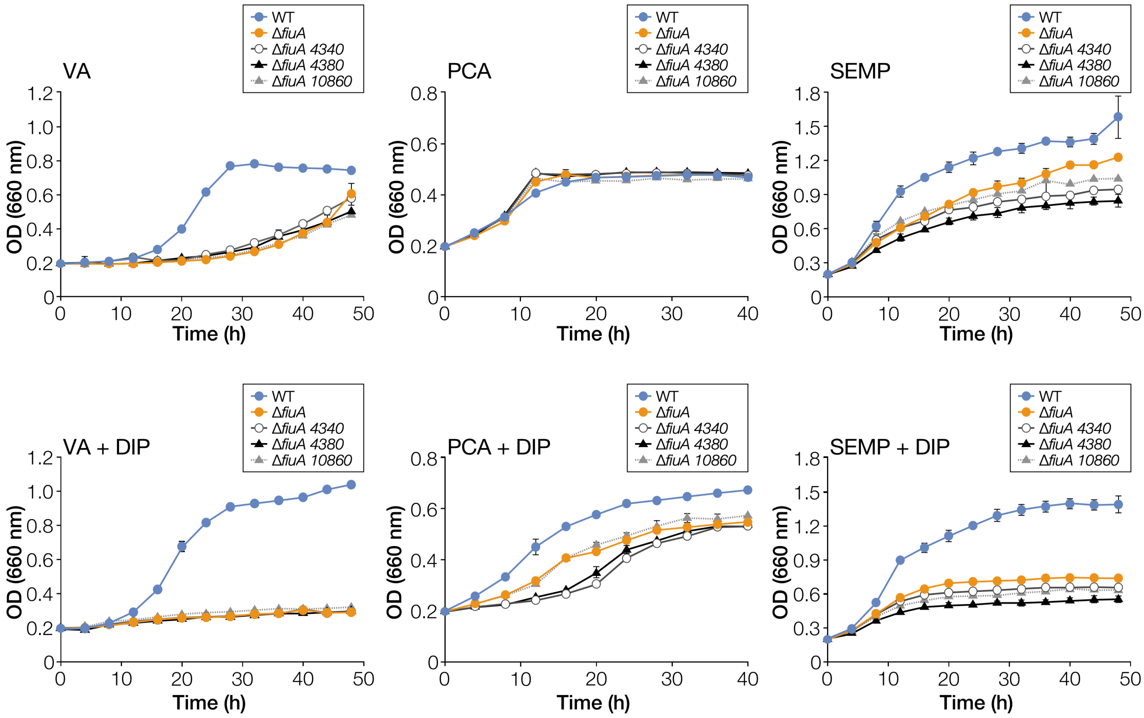


**Fig. S12. Growth of Δ*fiuA 4340*, Δ*fiuA 4380*, and Δ*fiuA 10860* on VA, PCA, and SEMP.** Cells of SYK-6, Δ*fiuA*, Δ*fiuA 4340*, Δ*fiuA 4380*, and Δ*fiuA 10860* were cultured in Wx medium containing 5 mM VA, 5 mM PCA, or SEMP in the presence or absence of 100 µM DIP. Cell growth was monitored by measuring the $OD_{660}$. Each value is the average ± the standard deviation of three independent experiments.

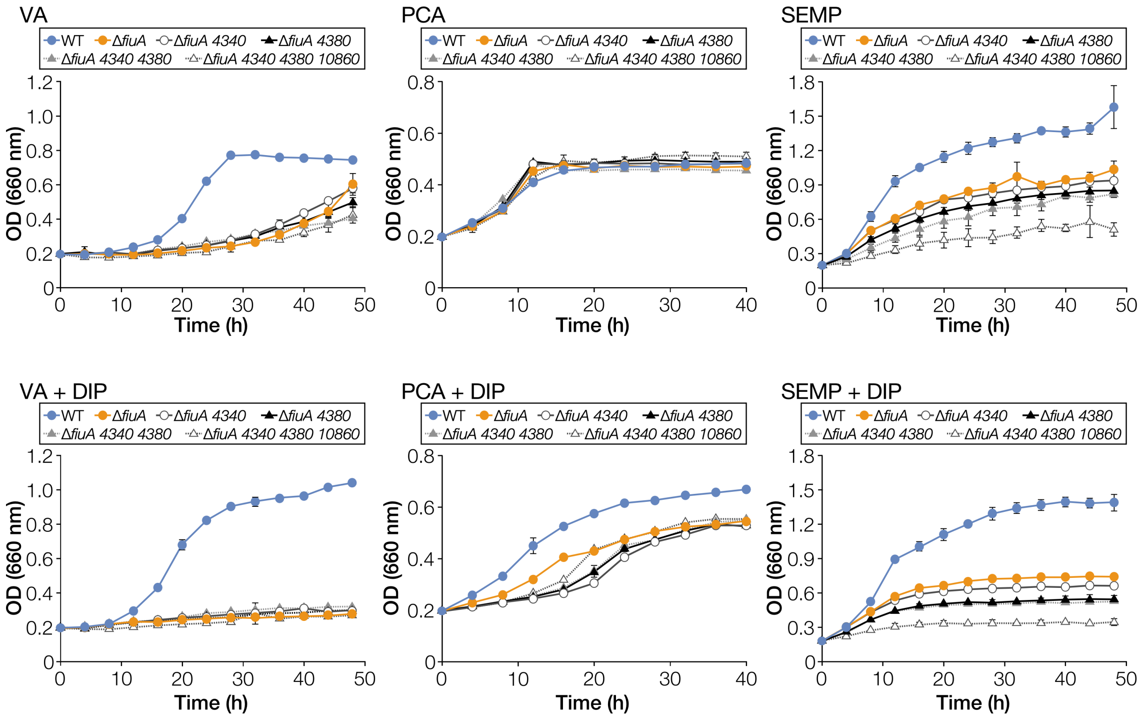


**Fig. S13. Growth of Δ*fiuA 4340 4380* and Δ*fiuA 4340 4380 10860* on VA, PCA, and SEMP.** Cells of SYK-6, Δ*fiuA*, Δ*fiuA 4340*, Δ*fiuA 4380*, Δ*fiuA 4340 4380*, and Δ*fiuA 4340 4380 10860* were cultured in Wx medium containing 5 mM VA, 5 mM PCA, or SEMP in the presence or absence of 100 μM DIP. Cell growth was monitored by measuring the $OD_{660}$. Each value is the average ± the standard deviation of three independent experiments.

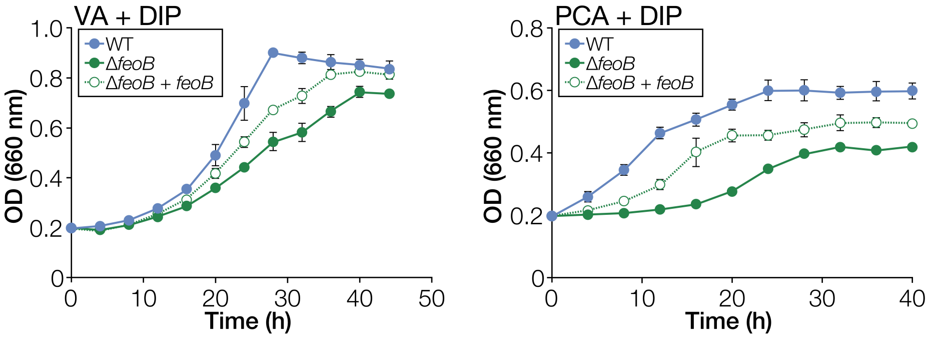


**Fig. S14. Growth complementation of Δ*feoB* under iron-limited conditions.** Cells of SYK-6(pJB861, vector), Δ*feoB*(pJB861), and Δ*feoB*(pJB-feoB) were cultured in Wx medium containing 5 mM VA or 5 mM PCA with 100 µM DIP and 1 mM *m*-toluate. Cell growth was monitored by measuring the $OD_{660}$. Each value is the average ± the standard deviation of three independent experiments.

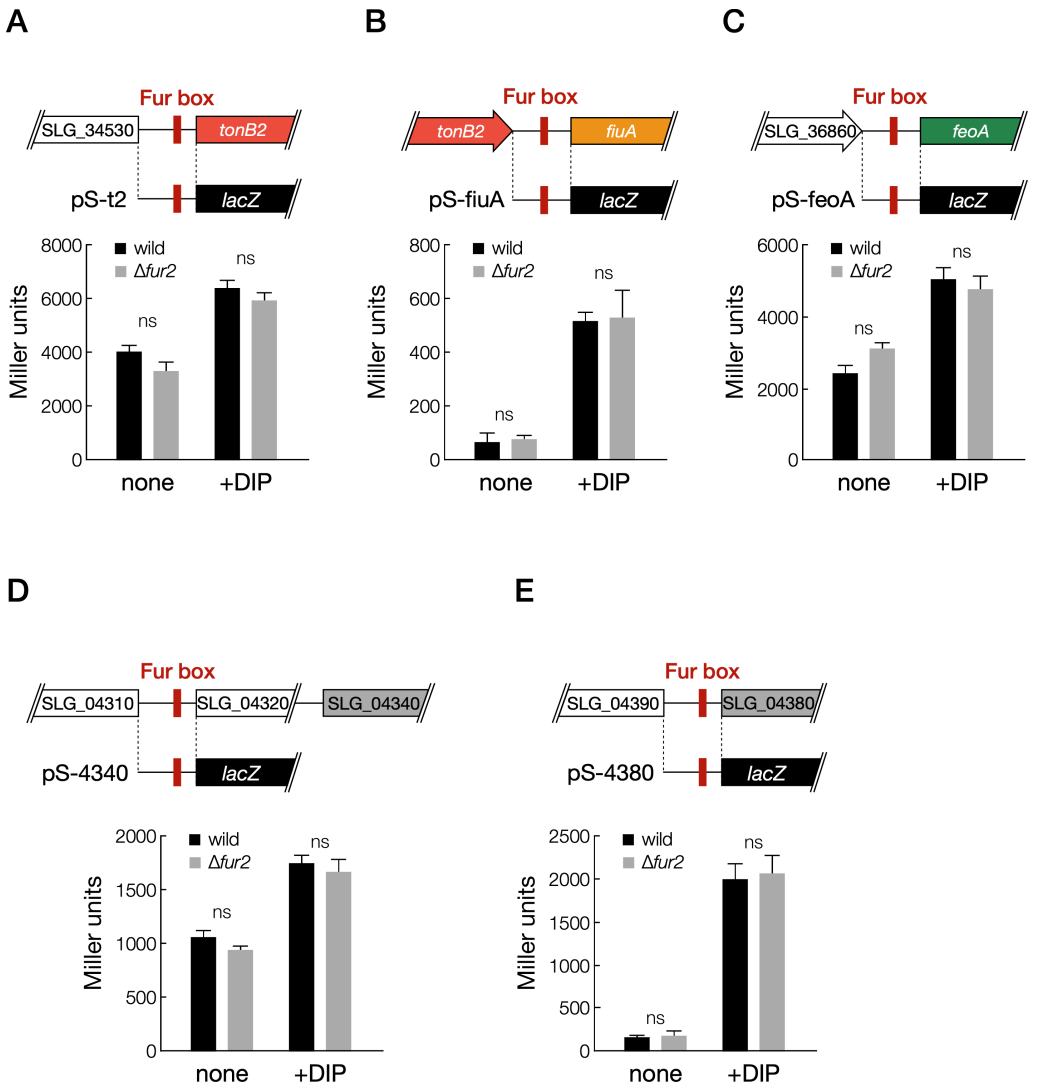


**Fig. S15. Disruption of *fur2* did not affect the promoter activities of iron transporter genes.** β-galactosidase activities of SYK-6 and Δ*fur2* cells harbouring pS-t2 (A), pS-fiuA (B), pS-feoA (C), pS-4340 (D), or pS-4380 (E) grown in Wx-SEMP with or without 100 μM DIP. The DNA fragments used for the promoter analysis are shown at the top. Each value is the average ± the standard deviation of three independent experiments. ns, $P > 0.05$ (two-tailed, unpaired *t*-test).

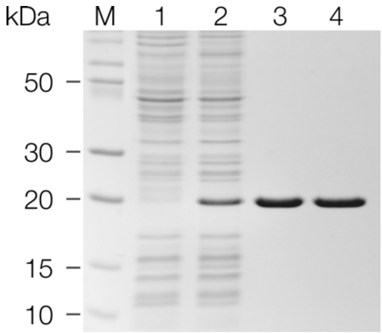


**Fig. S16. Purification of Fur1.** Proteins were separated using SDS-15% PAGE and stained with Coomassie Brilliant Blue. Lanes: 1, cell extracts of *E. coli* BL21(DE3) harbouring pET-16b (10 μg protein); 2, cell extracts of *E. coli* BL21(DE3) harbouring pET-fur1 (10 μg protein); 3, Fur1 purified by His Spin Trap (2.0 μg protein); 4, purified Fur1 after ultrafiltration (2.0 μg protein); M, molecular size markers.

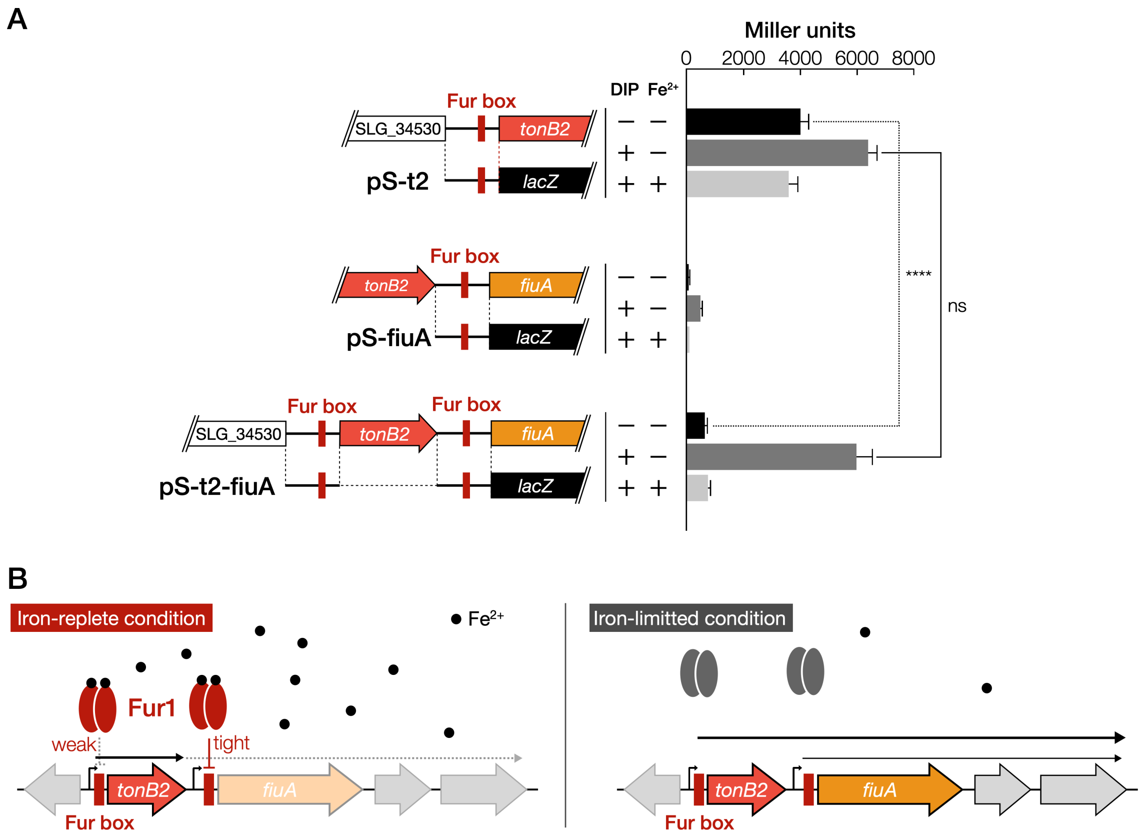


**Fig. S17. Transcription of *fiuA* is tightly regulated by Fur1.** (**A**) β-galactosidase activities of SYK-6 cells harbouring pS-t2, pS-fiuA, or pS-t2-fiuA grown in Wx-SEMP with or without 100 μM DIP and 100 μM $FeCl_2$. The DNA fragments used for the promoter analysis are shown on the left. Each value is the average ± the standard deviation of three independent experiments. ns, $P > 0.05$, ****, $P < 0.0001$ (two-tailed, unpaired *t*-test). (**B**) Proposed transcriptional regulation of the *tonB2-fiuA* operon. Under iron-replete conditions, transcription of *fiuA* from the *tonB2* promoter is interrupted by binding Fur1 to the Fur box of *fiuA* (left). Under iron-limited conditions, Fur1 is released from the Fur boxes, and *fiuA* is strongly transcribed from the *tonB2* promoter (right).

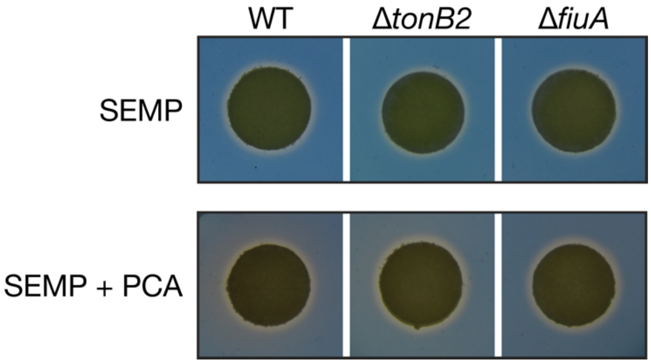


**Fig. S18. Production of siderophores by SYK-6, Δ*tonB2*, and Δ*fiuA*.** Cells of SYK-6, Δ*tonB2*, and Δ*fiuA* are incubated on Wx-SEMP-CAS agar plates with or without 1 mM PCA for 144 h.

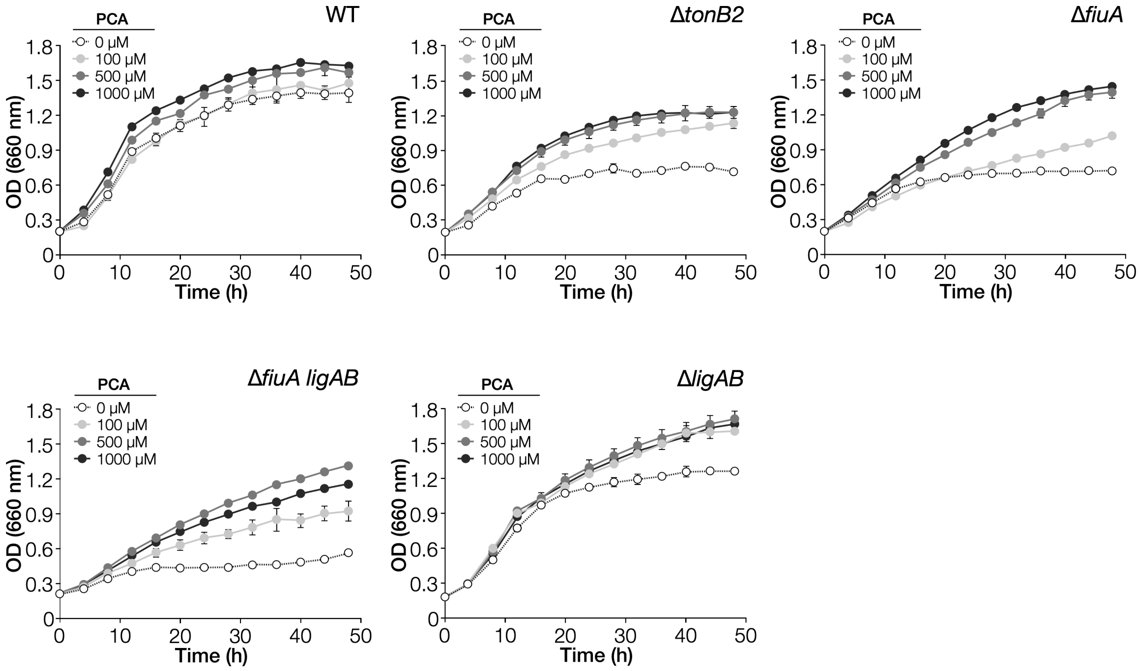


**Fig. S19. Effect of PCA addition on the growth of Δ*tonB2* and Δ*fiuA* on SEMP under iron-limited conditions.** Cells of SYK-6, Δ*tonB2*, Δ*fiuA*, Δ*fiuA ligAB*, and Δ*ligAB* were cultured in Wx-SEMP containing 100 μM DIP with or without PCA (100 μM, 500 μM, or 1000 μM). Cell growth was monitored by measuring the $OD_{660}$. Each value is the average ± the standard deviation of three independent experiments.

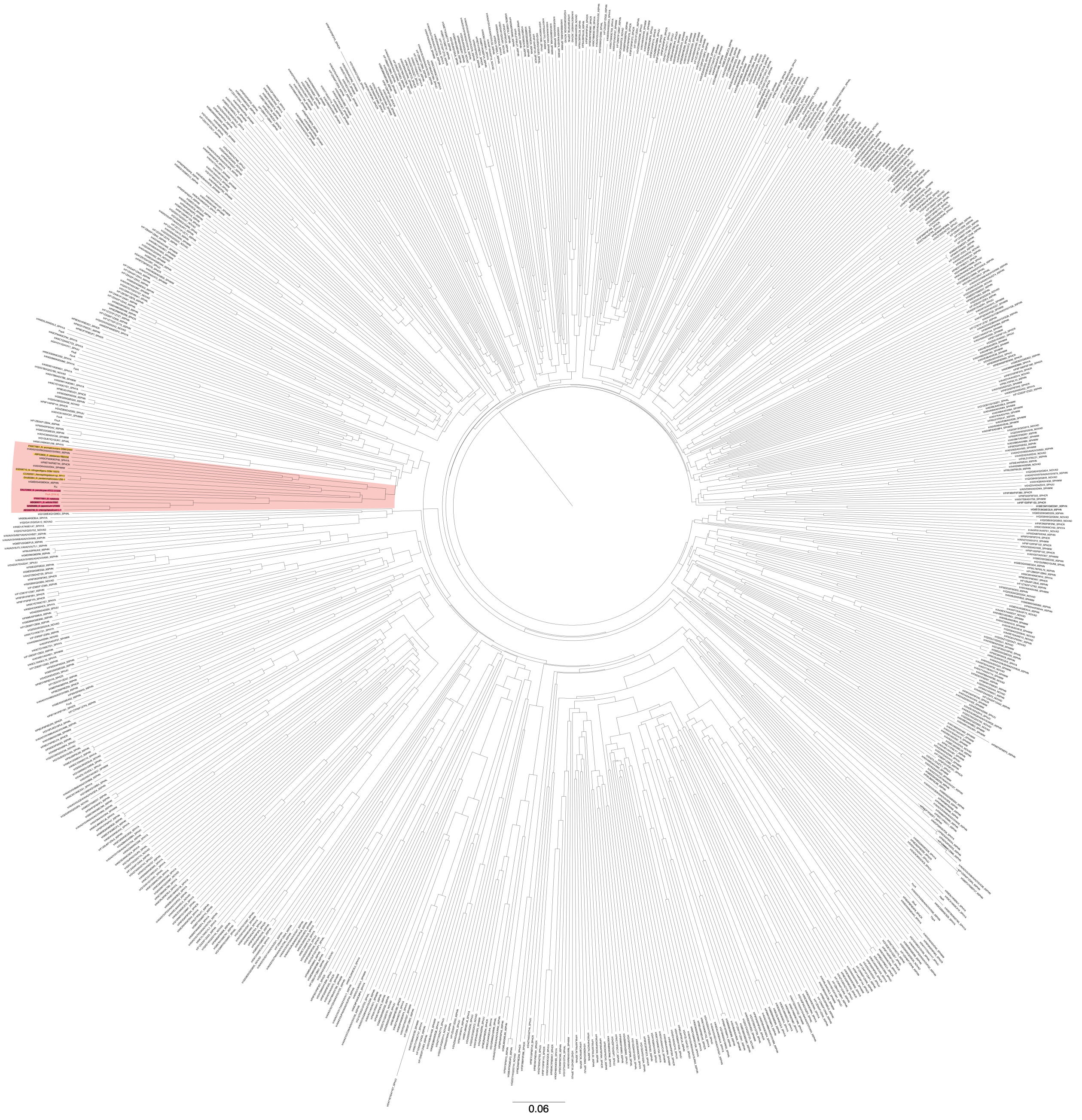

**Fig. S20. Phylogenetic tree of Sphingomonad TBDRs with known iron-uptake TBDRs.** TBDRs classified into the same clade with *fiuA* are highlighted with a red background. Among TBDRs listed in Table S3, TBDRs highlighted in magenta and yellow show >40% and 26–33% amino acid sequence identities with *fiuA*, respectively. The accession numbers of TBDR are shown in the figure, Table S2, and S3. The scale bar corresponds to 0.06 amino acid substitutions per position. A multiple alignment was performed using the Clustal Omega program[38].

**A**

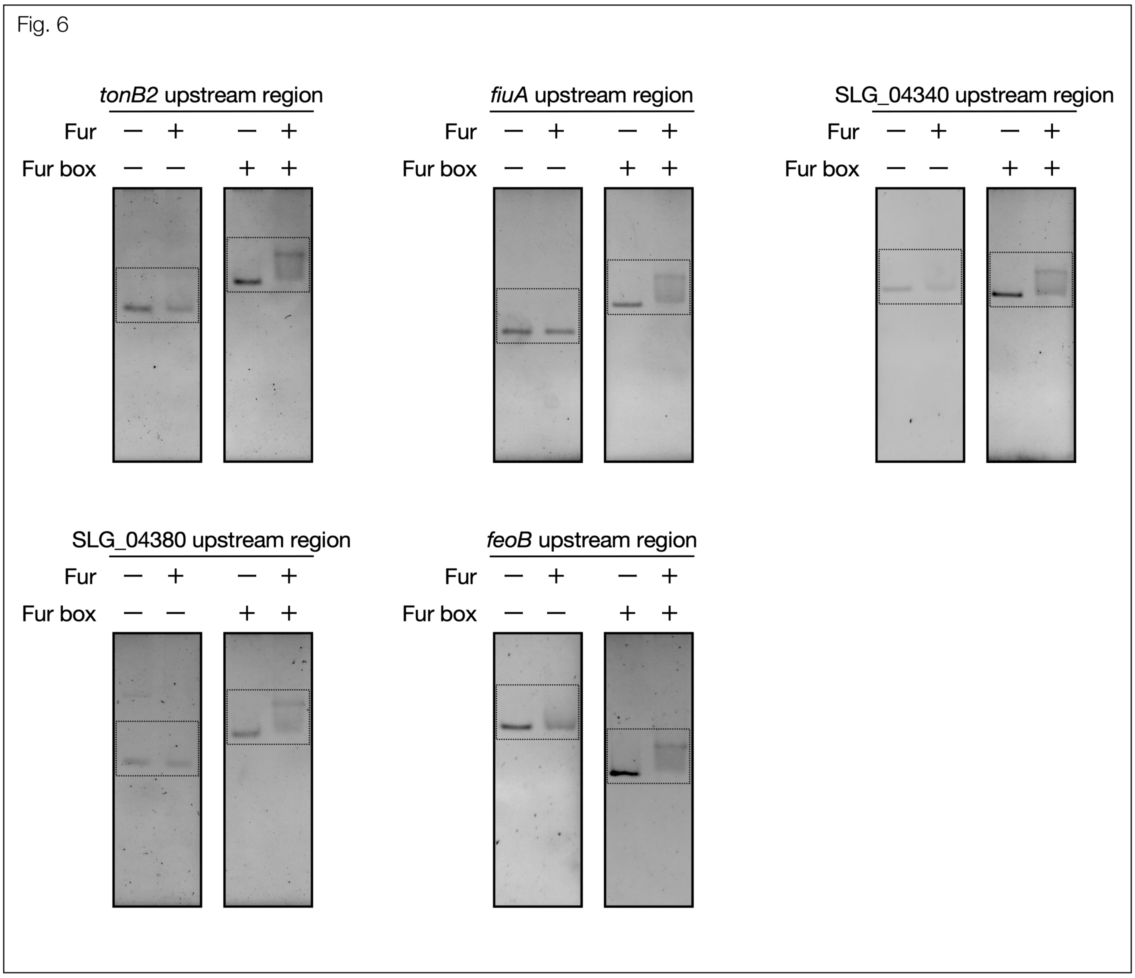


**B**

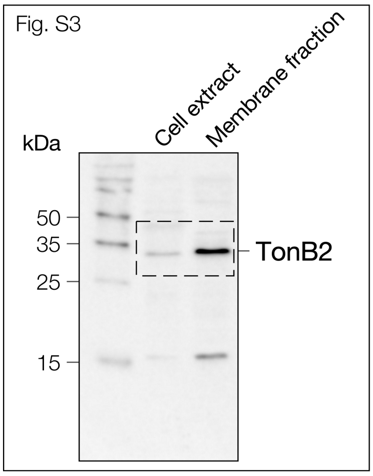


**Fig. S21. Uncropped western blots, ponceau staining, and EMSA images shown in Fig. 6, Fig. S3, Fig. S4, and Fig. S7.**

C

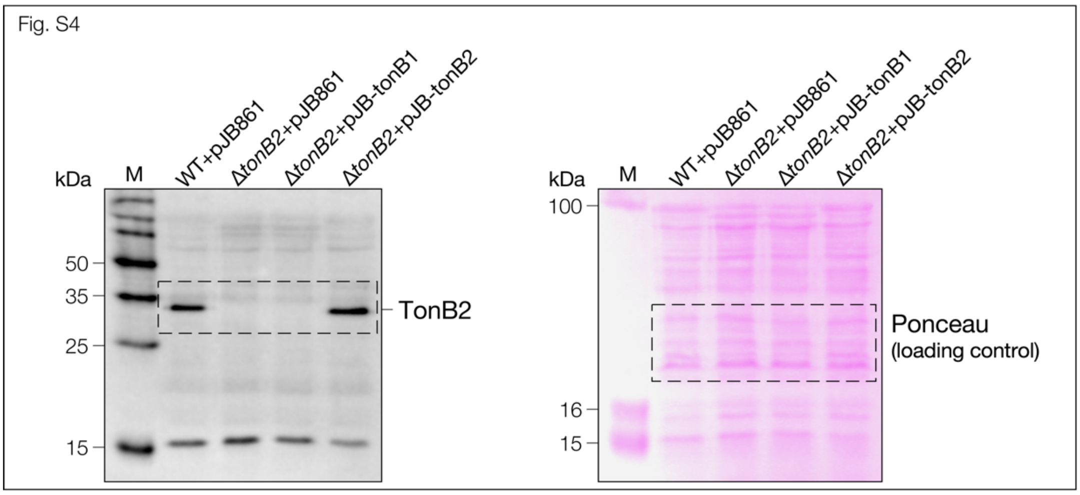


D

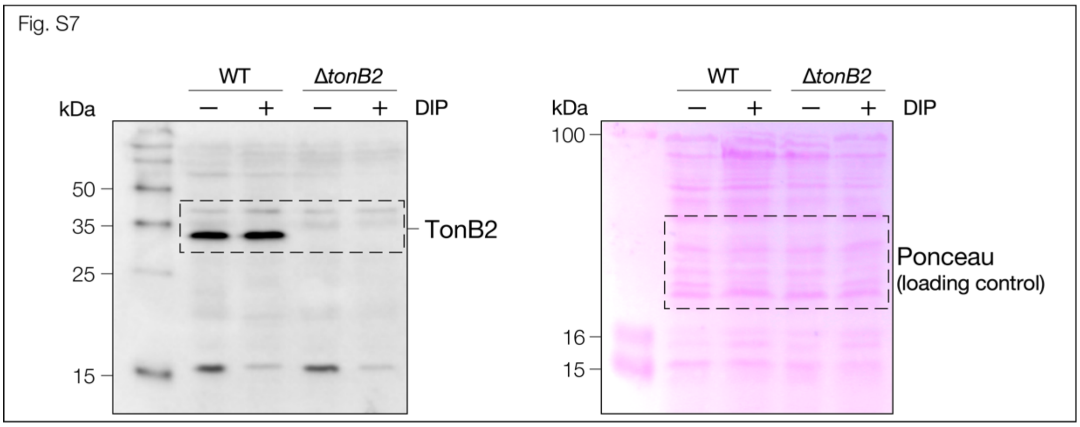


**Fig. S21.** −*continued.*
